# A novel fold for acyltransferase-3 (AT3) proteins provides a framework for transmembrane acyl-group transfer

**DOI:** 10.1101/2022.06.30.498268

**Authors:** Kahlan E. Newman, Sarah N. Tindall, Sophie L. Mader, Syma Khalid, Gavin H. Thomas, Marjan W. van der Woude

## Abstract

Acylation of diverse carbohydrates occurs across all domains of life and can be catalysed by proteins with a membrane bound acyltransferase-3 (AT3) domain (PF01757). In bacteria, these proteins are essential in processes including symbiosis, resistance to viruses and antimicrobials, and biosynthesis of antibiotics, yet their structure and mechanism is largely unknown. In this study, evolutionary co-variance analysis was used to build a computational model of the structure of a bacterial O-antigen modifying acetyltransferase, OafB. The resulting structure exhibited a novel fold for the AT3 domain, which molecular dynamics simulations demonstrated is stable in the membrane. The AT3 domain contains 10 transmembrane helices arranged to form a large cytoplasmic cavity lined by residues known to be essential for function. Further molecular dynamics simulations support a model where the acyl-coA donor spans the membrane through accessing a pore created by movement of an important loop capping the inner cavity, enabling OafB to present the acetyl group close to the likely catalytic resides on the extracytoplasmic surface. Limited but important interactions with the fused SGNH domain in OafB are identified and modelling suggests this domain is mobile and can both accept acyl-groups from the AT3 and then reach beyond the membrane to reach acceptor substrates. Together this new general model of AT3 function provides a framework for the development of inhibitors that could abrogate critical functions of bacterial pathogens.

## Introduction

Acyltransferase family 3 (*Acyl_transf_3*, AT3) domain (Interpro: IPR002656, PFAM: PF01757) containing proteins are found in all domains of life. These membrane-bound AT3 proteins are involved in acylation of a wide range of extracytoplasmic and surface bacterial polysaccharides (1), but are also important in Eukarya, for example in the regulation of lifespan in *Caenorhabditis elegans* (2) and in *Drosophila* development (3). In bacteria, where these proteins have been primarily studied, the resulting acylations have been shown to be involved in root nodulation (4), increase the efficacy of macrolide antibiotics (5), conferring resistance to lysozyme (6), influencing bacteriophage sensitivity (7), and altering antibody recognition (8–10). While AT3 domains most commonly exist as standalone proteins, there are many examples of AT3 domains fused with SGNH or alanine racemase domains, and fusions to other domains also exist in Eukarya (see Pfam architectures for PF10757). AT3 domains are predicted to have ten transmembrane helices (11–29); however, despite the wide-ranging functions of this family of proteins, there are currently no models for their overall structure and only limited information on mechanism.

The overall understanding of AT3 proteins is mainly derived from studies of these proteins in the context of bacterial virulence through changes in the cell surface (2,3). One example of this is the acetylation of lipopolysaccharide (LPS) and lipooligosaccharide (LOS) by AT3 domain containing proteins present in the inner (cytoplasmic) membrane of Gram-negative bacteria, including species of *Neisseria* (30), *Salmonella* (31,32), *Shigella* (33–38), *Haemophilus* (39,40), *Burkholderia* (41–44) and *Legionella* (27,45,46), that modify the O-antigen of the LPS during its synthesis and before LPS export to the outer membrane. Acetylation performed by these AT3 family proteins increases heterogeneity of the O-antigen, which can lead to an altered bacterial serotype and resistance to bacteriophage attack (31–36). Some of these reactions are carried out by AT3 only proteins, others by proteins with fused AT3 and SGNH domains. SGNH proteins are a family of proteins with hydrolase/transferase activity (47).

A second important process in bacteria that involves an AT3 protein is the modification of peptidoglycan that leads to resistance to both lysozyme and β-lactam antibiotics (29,48). One such protein, OatA, is an AT3 protein with attached SGNH domain (AT3-SGNH), that acetylates the MurNAc residue in peptidoglycan (29,48). Initially discovered in *Staphylococcus aureus* (29), OatA homologues have since been identified in many Gram-positive bacteria including *Listeria monocytogenes* (48), *Lactococcus lactis* (49,50) and *Streptococcus pneumoniae* (14,51,52). Furthermore, standalone AT3 proteins contribute an acyl group in the biosynthesis of macrolide antibiotics in *Streptomyces* species which increases antibiotic efficacy (5,53–55). These selected examples (see (1) for a recent review) illustrate that AT3 domain-containing proteins are involved in a range of highly relevant processes for bacterial pathogens and may be relevant for biotechnological applications.

Found in the cytoplasmic membrane, AT3 domains are highly hydrophobic integral membrane proteins predicted to contain 10 transmembrane helices (11–29). While the majority of AT3 domains consists of only an AT3 domain (standalone AT3), many are AT3 domains with an additional transmembrane helix (TMH) and C-terminal, periplasmic SGNH domain attached via a periplasmic linking region (AT3-SGNH) (56). Thanweer *et al*. (26) and Jones *et al*. (57) have performed PhoA-LacZɑ fusion analysis of the O-antigen acetyltransferase Oac from *Shigella flexneri* (a standalone AT3) and OatA from *Staphylococcus aureus* (an AT3-SGNH), respectively. Analysis by Thanweer *et al*. suggested that Oac has 10 TMH, with the N- and C-termini in the cytoplasm (26). This is consistent with fusion analysis of the O-antigen acetyltransferase OafB from *Salmonella* ser. Typhimurium (20). The *Salmonella* O-antigen acetyltransferases OafB and OafA are both AT3-SGNH fusion proteins and predicted to have an 11th TMH to allow the fused SGNH domain to be located in the periplasm (20,58). In contrast, Jones *et al*. proposed that OatA contain only 9 TMH with one large re-entrant loop (57); as an AT3-SGNH protein, OatA was also previously predicted to contain 11 TMH, orientating the SGNH domain in the periplasm (12,29). In addition, a large cytoplasmic loop was described between TMH 8-9 of OatA, where Oac has only short cytoplasmic loops but a longer periplasmic loop between TMH3 and 4 (26,59). Currently, this is the extent of the knowledge of the architecture of AT3 domains and there are no published experimentally determined protein structures.

AT3 proteins are predicted to transport acyl groups, including acetyl groups, from the cytoplasm across the cytoplasmic membrane to be transferred onto the receptor molecule (6). However, the mechanism is currently largely unknown. The acetyl group has been proposed to be donated by acetyl coenzyme A (acetyl-CoA)(24). Three arginine residues on the cytoplasmic side of TMH1 and 3 have been identified as highly conserved in the PFAM HMM logo, and in both standalone AT3 and AT3-SGNH proteins these residues are essential for function (26,58). Arginine residues have been shown to be involved in binding of the 3’-phosphate group of acetyl-CoA (60) suggesting that these conserved residues may interact with the proposed acetyl group donor. Previous *in vitro* studies suggest that standalone AT3 proteins CmmA, MdmB and Asm19 are able to utilise acetyl-CoA as an acetyl donor (5,61,62). Importantly, Jones et al. (57) found OatA from *Staphylococcus aureus* (an AT3-SGNH protein) is able to hydrolyse the acetyl group from acetyl-CoA and transfer it to the a peptidoglycan-like acceptor substrate, effectively observing the entire catalytic cycle *in vitro* (57). Thus, the available data indicate that acetyl-CoA is the most likely acetyl group donor is acetyl-CoA, but how this interacts with the AT3 domain or how the acetyl group can cross the membrane, is as yet unknown.

In addition, site-directed mutagenesis identified three conserved tyrosine residues which are required for function (57). One of these Tyr residues is predicted to be located in the periplasm and is proposed to be involved in transfer of the acyl group from the AT3 domain to the SGNH domain in AT3-SGNH proteins (57). The catalytic triad of the SGNH domain of OafB from *Salmonella enterica* subsp. *enterica* serovar Typhimurium is known to be required for acetylation of the rhamnose moiety of the O-antigen of LPS (58), suggesting that in standalone AT3 proteins an equivalent, as yet unknown, partner protein may exist. If this were the case, it would suggest that the system may be similar to that of the PatA and PatB peptidoglycan acetyltransferase system. In this system it is hypothesised that the MBOAT (membrane-bound O-acyltransferase) protein PatA transports the acetyl group across the membrane where it is transferred to the peptidoglycan substrate by PatB (an SGNH domain containing protein) (63).

To further study the AT3-SGNH family of proteins, OafB from *Salmonella enterica* subsp. *enterica* was used as a model system. OafB acetylates the rhamnose moiety of the O-antigen repeating unit of LPS. Modification of the O-antigen has repeatedly been identified as important for virulence and persistence of bacteria (27,30,39,45). Similarly, O-acetylation by OafB has been acknowledged in the development of vaccines against *Salmonella* ser. Paratyphi A, where acetyl groups were shown to be required to elicit bactericidal antibodies (64,65). Furthermore, OafB-mediated acetylation in invasive non-typhoidal Salmonella (iNTS) alters susceptibility to bacteriophage (20). OafB consists of an AT3 domain linked to an SGNH domain via an 11th TMH and periplasmic linking region (AT3-SGNH) (58) (**Fig. 1A**). The structure of the SGNH domain and periplasmic linking region was previously solved using x-ray crystallography (58). The structure was found to be similar to that of other SGNH domains, but the periplasmic linking region formed a structured extension suggesting the AT3 and SGNH domains are likely to exist in close proximity and may even interact (58). Herein we use RaptorX to determine a computationally derived structure for the transmembrane region, including the 10-TMH AT3 domain, of OafB. We show this novel structure to be stable under physiological conditions *via* molecular dynamics simulations. This model is integrated with the existing x-ray structure of the SGNH domain and published mutational data to allow additional structure-function analysis. Utilising both classical molecular dynamics simulations and quantum mechanical calculations, we consider the acetyl donor and acceptor substrates to generate a refined functional model for this important family of membrane proteins.

**Figure 1:**
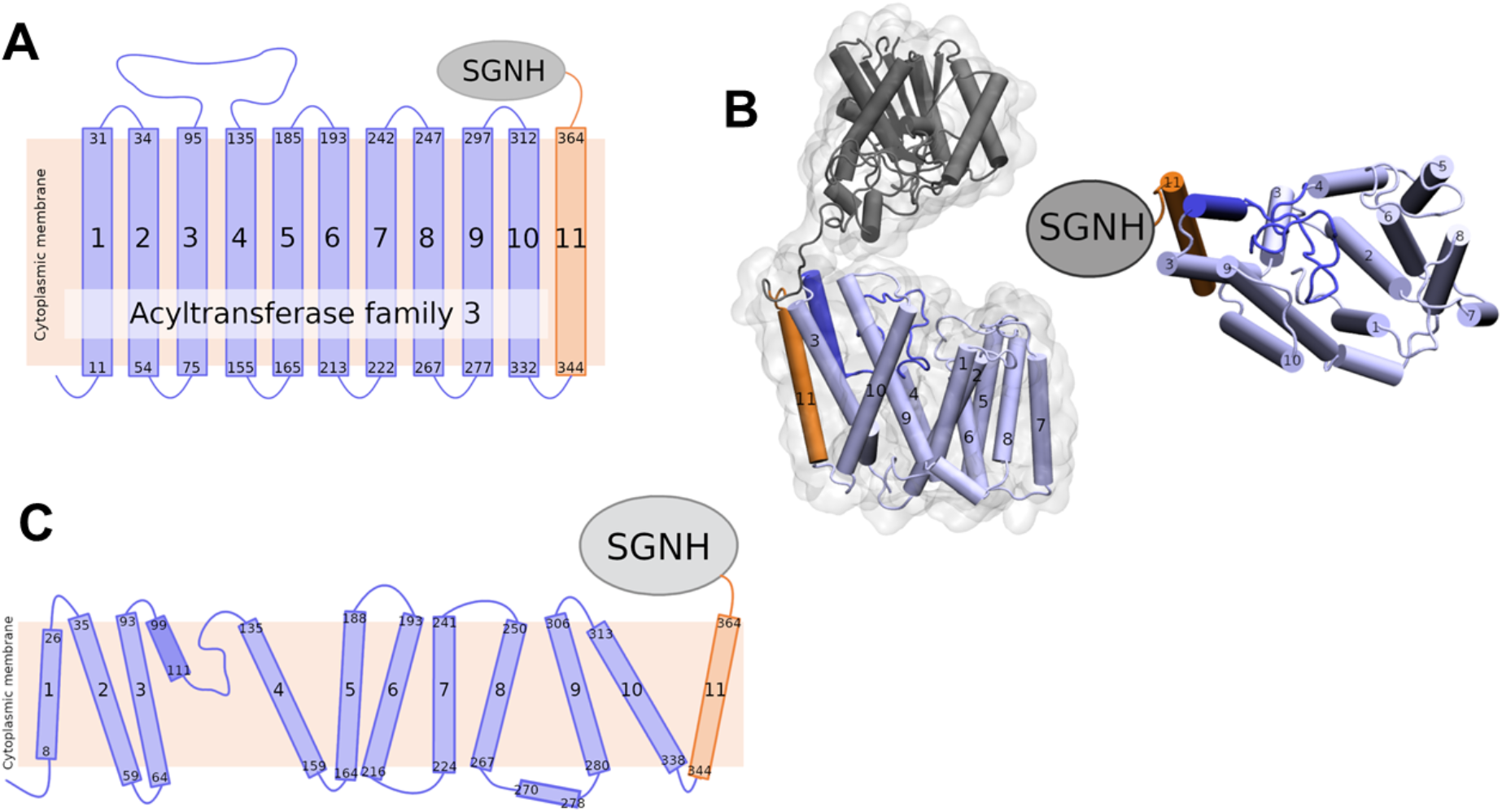
Topology and predicted structure of OafB. **(A)** TOPCONS topology prediction of OafB with N- and C-termini of each TMH indicated. Consistent with the RaptorX structure, TOPCONS predicts short cytoplasmic loops and a long periplasmic loop between TMH 3 and 4. **(B)** RaptorX predicted structure of OafB with AT3 domain and TMH 11 coloured light blue, periplasmic loop 3-4 coloured blue, SGNH domain coloured grey with the periplasmic linking region in orange and additional helix in teal. **(C)** Topology schematic of OafB based on the RaptorX predicted structure. The RaptorX structure has 11 TMH, with a long periplasmic loop between TMH 3 and 4 consisting of a short helix followed by an unstructured region. TMH1-10 form the AT3 domain and are coloured blue, TMH11 forms the linking region between the AT3 and SGNH domains and is coloured orange.

## Results & Discussion

### RaptorX model of OafB supports specific topology predictions and identifies new features

The structure of OafB was predicted by RaptorX (66–68) using the protein sequence of a *Salmonella* rhamose O-acetyltransferase (OafB). This method has been successfully validated with numerous proteins (69–73). The structure of OafB consists of two key domains: the AT3 domain and SGNH domain (**Fig. 1B**). The transmembrane region has 13 helices, of which 11 completely span the membrane (**Fig. 1B, C**). The AT3 domain (residues 1-338) has 10 TMH and an additional 11th TMH (residues 339-376) fused to the periplasmic linking region (containing SGNH-ext, residues 377-421), which facilitates a periplasmic location of the fused SGNH domain (residues 422-640) (58), as predicted from simple TOPCONS topological analysis (74,75). The topological analysis also predicted a loop region between helices 3 and 4 (**Fig. 1A**); in the RaptorX model this region is clearly suggested to be a short re-entrant loop consisting of a short, well-structured helix (residue 99-111) followed by a ∼24 residue-long unstructured region **(Fig. 1B,C**). This means that there are no long hydrophilic loops in the structure on either side of the membrane (58).

The structure of the SGNH domain and periplasmic linking region of OafB was previously solved using x-ray crystallography (58). The RaptorX structure of the same region showed remarkable similarity, with a root-mean-square deviation (RMSD) of 2.8 Å. As seen in the crystal structure, the SGNH domain from OafB resembles a typical SGNH domain but with an additional helix and structured extension. Neither of these regions are seen in the structures of other SGNH domains in the RCSB PDB; despite this, RaptorX predicts these regions with high accuracy. There are very few residues between the C-terminus of TMH11 (residue 367) and the N-terminus of the structured extension (residue 377), suggesting that the AT3 and SGNH domains are likely in close proximity and may interact (58). However, in the RaptorX structure of OafB, the two domains show limited interaction and are not in close proximity (**Fig. 1B**).

Similar to RaptorX, AlphaFold (76) predicted 11 transmembrane helices, with the SGNH domain in the periplasm. The structures predicted for residues 1-338 (TMH1-10 of the transmembrane domain, the AT3 domain) showed 97% coverage (RMSD of 3.04 Å) and a template matching score of 0.85, indicating the same structure across the two models (**Supp. Fig. S1**).

### Diverse bacterial AT3 domain-containing proteins share a common 10 TMH structure

Having built a model for our primary experimentally characterised protein, we expanded the analysis to include other important AT3 domain-containing proteins, including both additional AT3-SGNH fusion proteins and standalone AT3 proteins. The RaptorX generated structures of the AT3-SGNH proteins OatA from *Staphylococcus aureus* (OatA-SA), OatA from *Listeria monocytogenes* (OatA-LM) and PglI from *Neisseria gonorrhoeae* (PglI-NG) all closely resembled the OafB (**Fig. 2A**), with an RMSD of less than 3 Å for the AT3-domain over the 10 TMHs. Importantly, as they all contain a C-terminal SGNH domain that functions extra-cytoplasmically, they all contain an 11th TMH to enable correct localisation of the fused domain (**Fig 2A**) (23,58,77,78). In addition, the core of the SGNH domains are also similar (79) with variation occurring in the length of the linking regions that connect the AT3 to the SGNH. While the PglI-NG protein closely resembles OafB with a short, structured linker (1), OatA proteins show a more extended, and potentially more mobile and flexible, structure, locating the SGNH domain further from the AT3 domain (**Fig. 2A**).

**Figure 2:**
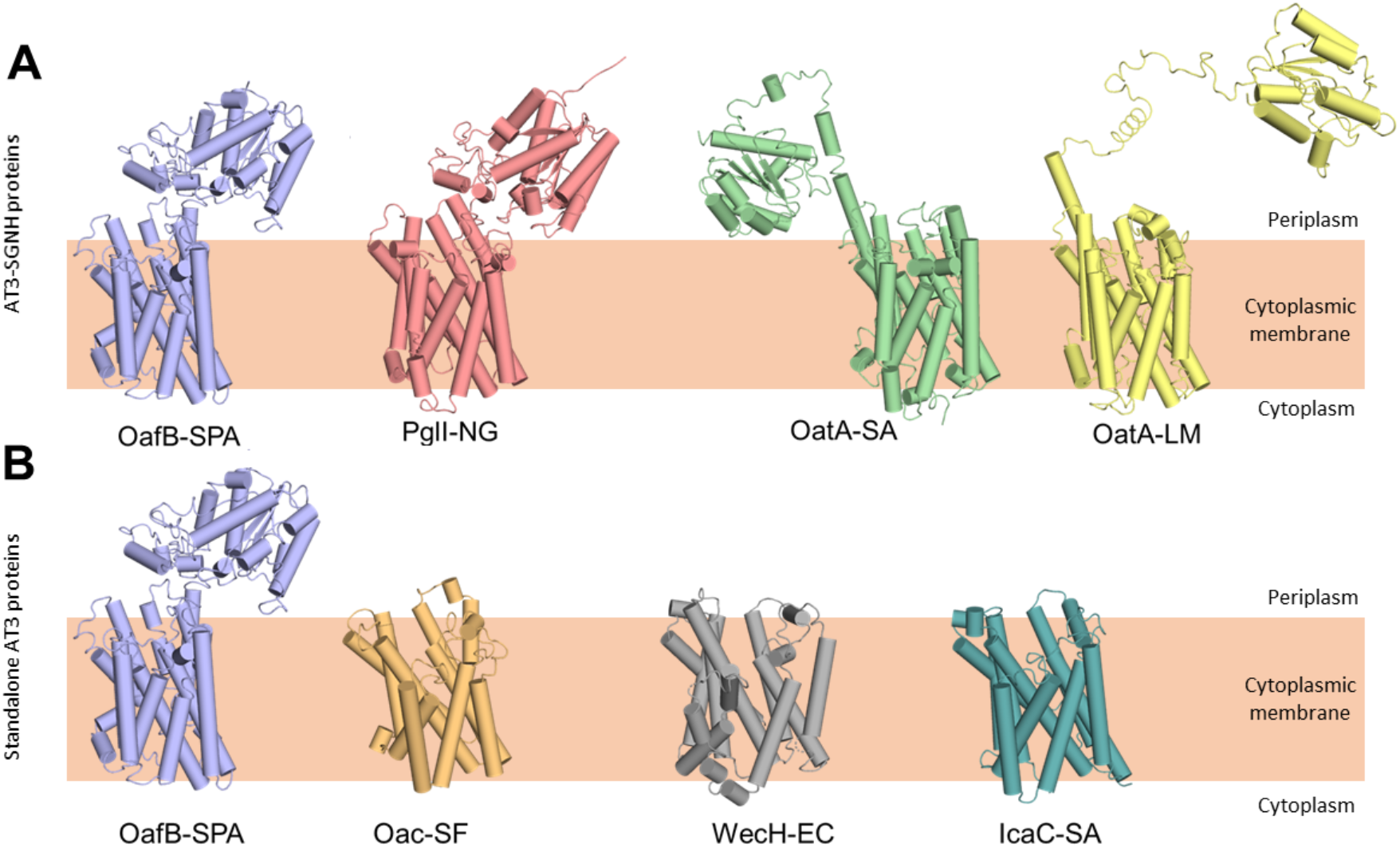
RaptorX predicted structures of AT3 domain containing proteins. **(A)** Fused AT3-SGNH proteins with OafB for comparison. Left to right: OafB from *Salmonella enterica* subsp. *enterica* ser. Paratyphi A (OafB); PglI from *Neisseria gonorrhoeae* (PglI-NG); OatA from *Staphylococcus aureus* (OatA-SA); OatA from *Listeria monocytogenes* (OatA-LM). **(B)** Standalone AT3 proteins with OafB for comparison. Left to right: OafB-SPA; Oac from *Shigella flexneri* (Oac-SF); WecH from *Escherichia coli* (WecH-EC); IcaC from *Staphylococcus aureus* (IcaC-SA). The structure of the AT3 domain is largely conserved between these proteins.

Expanding the analysis to standalone AT3 proteins, the RaptorX outputs from Oac from *Shigella flexneri* (Oac-SF), WecH from *Escherichia coli* (WecH-EC) and IcaC from *Staphylococcus aureus* (IcaC-SA) were analysed (**Fig. 2B**). While the RMSD of these compared to the AT3-domain of OafB was more variable (RMSD of 4.5 - 6.7 Å when compared to residues 1-376 of OafB) the overall 10 TMH features were conserved and are consistent with the experimentally-determined topology of Oac-SF (10,26). However, a recent analysis of *Staphylococcus aureus* OatA combining LacZ-PhoA fusion data with *in silico* predictions (56), led to an alternative model for the topology of the AT3 domain with 9 TMH, a further re-entrant helix and a long cytoplasmic loop preceding the final TMH that leads to the SGNH domain in the periplasm. This topology is not seen in any of our models, including similar models generated in AlphaFold (data not shown) and these conflicts will be discussed later.

### The AT3 domain defines a novel and stable membrane protein fold

Using our new structural models for AT3 proteins, we assessed their similarity to known structures using the DALI server (80). The closest structural homologues consisted of small helical bundles that were fragments of larger proteins and no significant matches to full length proteins were identified. Looking specifically at other membrane proteins with 10-11 TMH (3M73, 4J72, 4KJR, 1RH5) we could not see similarities with the AT3 structure (RMSDs between 12-24 Å) and the arrangement of TMH is very different. Finally, we compared our AT3 structure to that of the other known family of membrane-bound acyltransferases, the MBOAT proteins. These also function in the acylation of complex extracytoplasmic carbohydrates (81) and previous analysis has shown an MBOAT protein to be functionally interchangeable with a standalone AT3 protein (19,82) (**Fig. 3B**), again suggesting that these proteins have closely related functions. The structure of DltB (an MBOAT protein from *Streptococcus thermophilus*) (81) similarly contains 11 TMH, however, the helical arrangement is entirely different (**Fig. 3C,D**). Consistent with a core 10 TMH fold in the AT3-SGNH protein, the 11th TMH present sits on the outside of the bundle as would be expected for a non-essential feature of the AT3 domain, in contrast to the 11th TMH in DltB (**Fig. 3C**).

**Figure 3:**
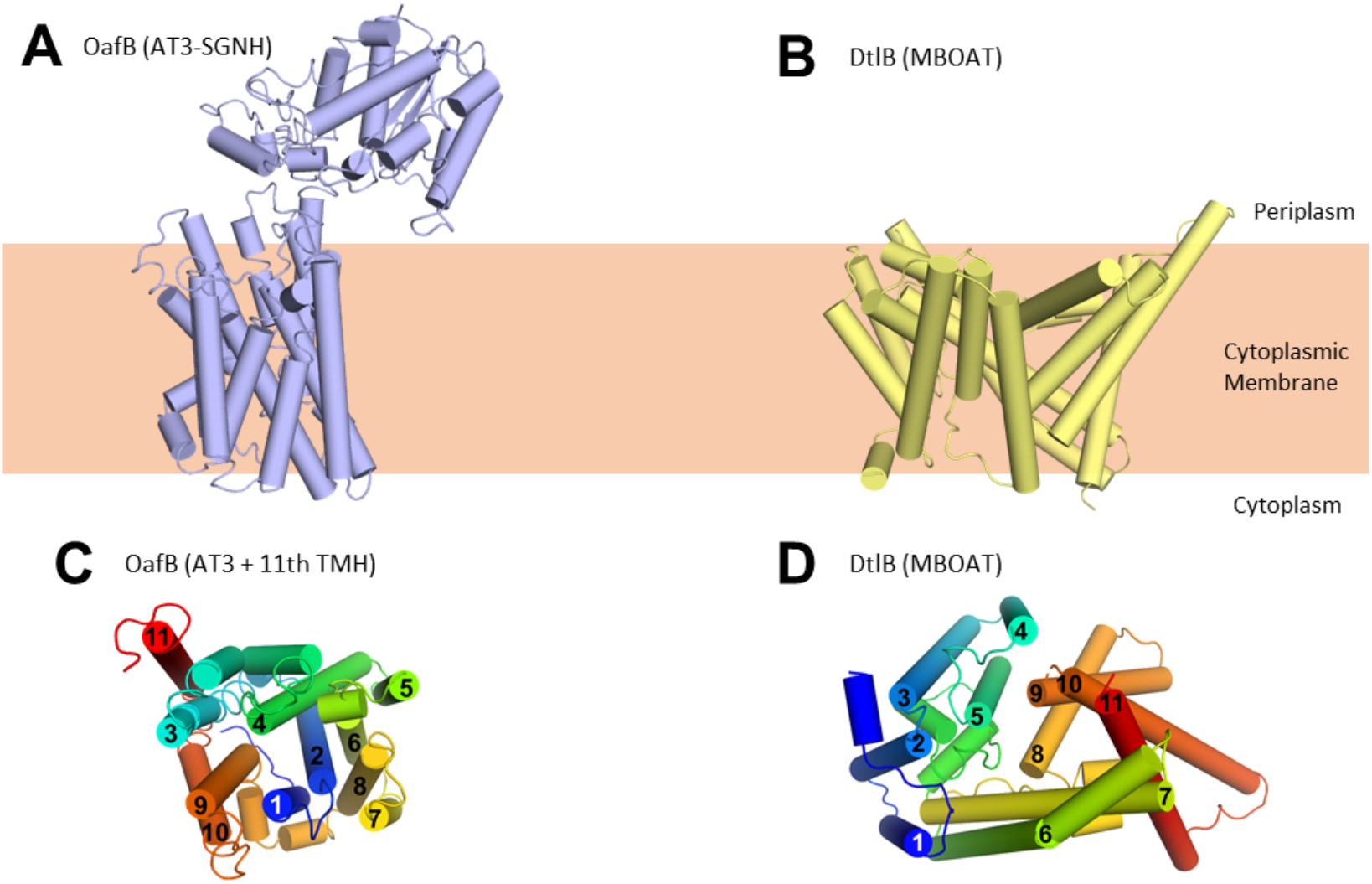
Structure of OafB, an AT3-SGNH protein (left, panels **A** and **C**), compared to DltB from *Streptococcus thermophilus*, an MBOAT (Membrane Bound O-Acyltransferase) protein (right, panels **B** and **D**). Both OafB and DltB have 11 TMH **(C, D)** however the arrangements differ considerably. **(C)** OafB and **(D)** DltB viewed from periplasm, coloured blue to red, N-to C-termini with transmembrane helices numbered.

To investigate the biological plausibility of the proposed AT3 fold, we applied Molecular Dynamics (MD) techniques to emulate the behaviour of the protein in a physiological environment, and to extract meaningful data on the dynamics and structure-function relationships of the system. The RaptorX model of the transmembrane domain (TMH1-11 of OafB, residues 1-376) was embedded in a model *Escherichia coli* inner membrane in 150 mM KCl solution. When subjected to MD simulations of 50 and 100 ns at 320 and 303 K, respectively, the protein model remained stable within the membrane. There was no significant unfolding observed; secondary structure analysis indicates that the alpha-helical segments of the transmembrane domain are maintained throughout and the RMSD of the protein backbone remained below 0.5 nm at both temperatures (**Supp. Fig. S2A**). The root-mean-square fluctuation (RMSF) by residue, as expected, is highest for the termini and for residues in unstructured loops, and lower for the transmembrane helices (**Supp. Fig. S2B)**. Together these data suggest that the computational approach described a novel, stable membrane protein fold for AT3 proteins.

### The AT3 domain from OafB has a cavity lined with essential residues forming a putative acetyl-CoA binding site

Having established that the AT3 domain constitutes a novel stable fold in the membrane we then wished to use this structural model to try and understand the mechanism of this membrane-bound enzyme. A large cavity with a volume of ∼3,620 Å^3^ can be observed in the AT3 protein on the cytoplasmic side, which is enclosed by enclosed by TMH 1, 2, 9, and the unstructured loop between TMH 5 and 6 (**Fig. 4A**). The top end of this cavity is blocked by the novel re-entrant loop between THM 3-4, which in contrast to TOPCONS predictions (**Fig. 1A**) is not in the periplasm but resides within the transmembrane domain (**Fig. 4A**). Examination of the positioning of core amino acids known to be important and/or essential for function of different AT3 proteins onto the predicted structure reveals that many of these in fact line this cytoplasmic cavity in the protein (**Fig. 4C**).

**Figure 4:**
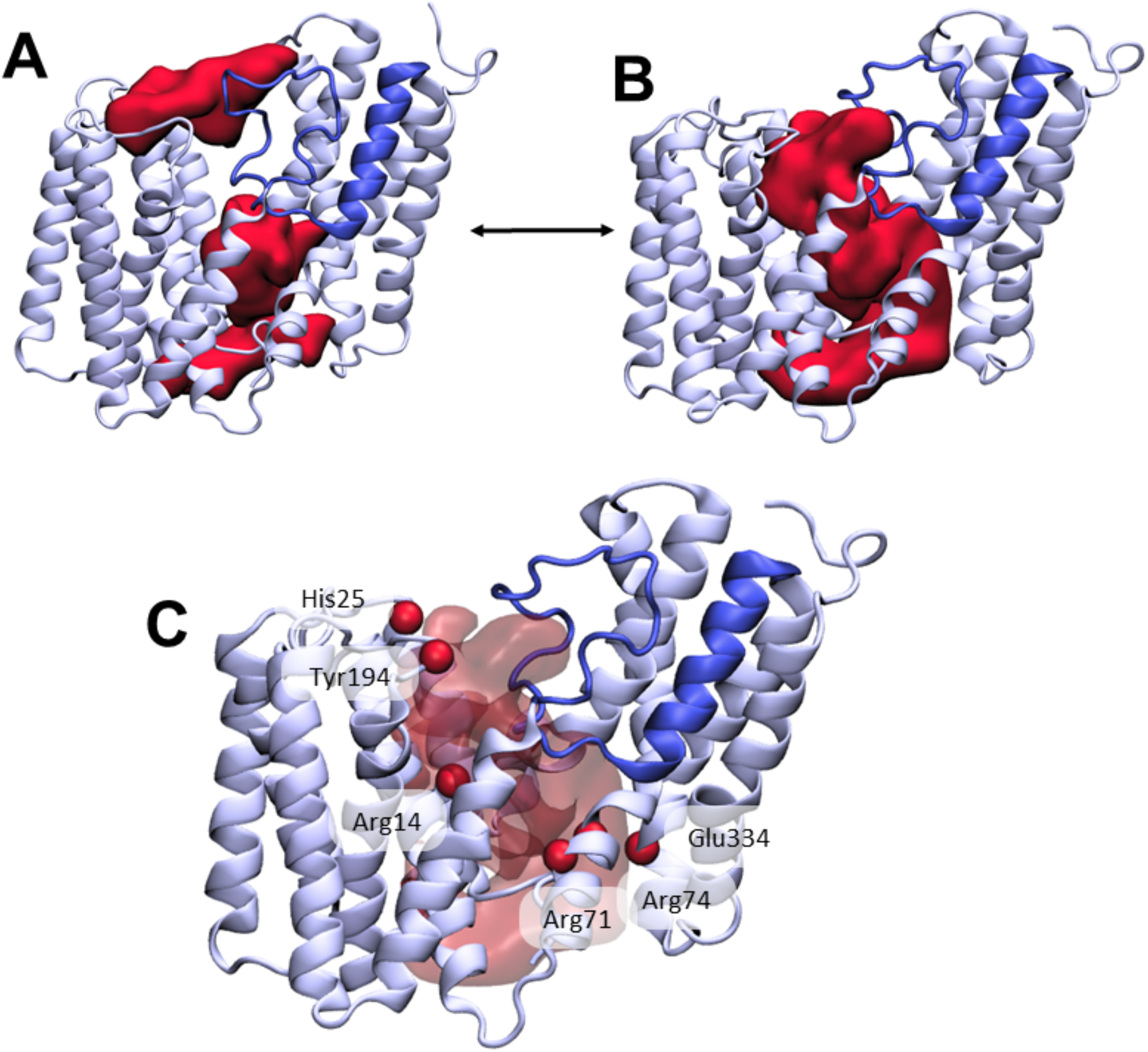
The loop between TMH3 and 4 controls the formation of a transmembrane channel lined with essential residues. **(A)** Initial equilibrated structure of the transmembrane domain (residues 1-376) of OafB. Residues 1-94 and 136-376 in pale blue, residues 95-135 in blue. The two largest cavities identified by the ProPores2 server are shown as red surfaces. Initially, the loop between TMH3 and 4 occludes the central pore in the AT3 domain. **(B)** Structure of the transmembrane domain after 50 ns equilibrium MD at 320 K. The loop between TMH3 and 4 is dynamic – this snapshot from the end of the simulation shows the loop to have moved away from the center of the cavity to allow a channel to form. This channel could be occupied by the acetyl group donor. **(C)** Same snapshot as (B), but pore shown as a transparent red surface. Important conserved residues shown as red spheres: these residues line the pore in the center of the AT3 domain.

Two conserved motifs have been identified in AT3 domains: an R-X_10_-H motif located in TMH1 (Arg14 and His25 in OafB, **Fig. 4C**) (58); and an RXXR motif located on the cytoplasmic face of TMH3 (Arg71 and Arg74) (17,20,24,83). The four residues found in these motifs have all been identified in experimental studies as being essential for function (20,25,57,58,84). Both arginine and histidine residues have previously been implicated in other acetyltransferase proteins as important for binding of acetyl-CoA (60,81,85). Perhaps significantly, all three Arg residues are in close proximity in positions approximately in the middle of the membrane, a usually unfavourable location, suggesting a key function in the mechanism of the enzyme.

In addition, Jones et al. performed site-directed mutagenesis on a number of residues in OatA from *Staphylococcus aureus* (57). These identified, amongst others, three conserved tyrosine residues which are required for function (57). It is thought that Tyr206, located in the periplasm (equivalent residue in OafB Tyr194, **Fig. 4A**), is involved in transfer of the acetyl group from the AT3 domain to the SGNH domain (57). Similarly, a catalytic triad consisting of this aforesaid tyrosine, plus a glutamic acid and a histidine in the AT3 domain was proposed to remove the acetyl group from acetyl-CoA (57), allowing transport across the membrane (the three equivalent residues are highlighted in **Figure 4A** – Tyr194, His25 and Glu334). In the RaptorX structure of OafB, Tyr194 and His25 are in close proximity (C_α_ separated by 8 Å) and both sit on the periplasmic face of the membrane. However, Glu334 is located on the cytoplasmic side very distant from His25 and Tyr194, suggesting that it is not part of a catalytic triad.

Together this mapping suggests two important sites in the protein, a mid-membrane region of positive charge created by 3 conserved Arg residues and a potential catalytic site on the periplasmic surface comprising at least His25 and Tyr194. In the RaptorX model these two features of the protein are physically separated by the loop 3-4 region (**Fig. 4A**). During equilibrium MD simulations this loop is in fact dynamic (**Fig. 4B**). Structures taken from later in the simulations indicate that the loop can move outwards (displacement of residues on the order of 3-4 Å) from the centre of the cavity towards TMH2, allowing the potential formation of a pore between the cytoplasmic and periplasmic surfaces of the transmembrane domain, which could be a critical stage in the mechanism of the acyltransferase in the trans-membrane movement of acyl-groups (**Fig. 4B**).

### The AT3 domain facilitates presentation of acyl-CoA molecules to the periplasm

Any model for an acyl-coA dependent acylation of an extracytoplasmic acceptor sugar requires the transfer of the cytoplasmically-located acyl group across the membrane. The inner cavity and dynamic conversion of this to a pore, could provide the route for acyl-CoA molecules to enter the protein in a way to present the acyl group for use in the catalytic process (**Fig. 5A, left**). OafB would use acetyl-CoA as the substrate, which is an extended molecule with a volume of ∼630 Å^3^. To test this hypothesis, one molecule of acetyl-CoA was pulled into this channel (entrance delimited by the N-terminus and TMH9 and 10) from the cytoplasmic entrance of the AT3 domain of a hybrid full-length OafB model (assembly discussed in the following section) using steered MD (SMD). It was found that this channel could indeed accommodate acetyl-CoA (**Fig. 5A, right**) and, significantly, the thioester bond was positioned close to the essential His25 residue within the membrane. The conserved Arg14 was also positioned to help coordinate the 3’-phosphate of acetyl-CoA, as previously predicted (58) (**Fig. 5A**).

**Figure 5:**
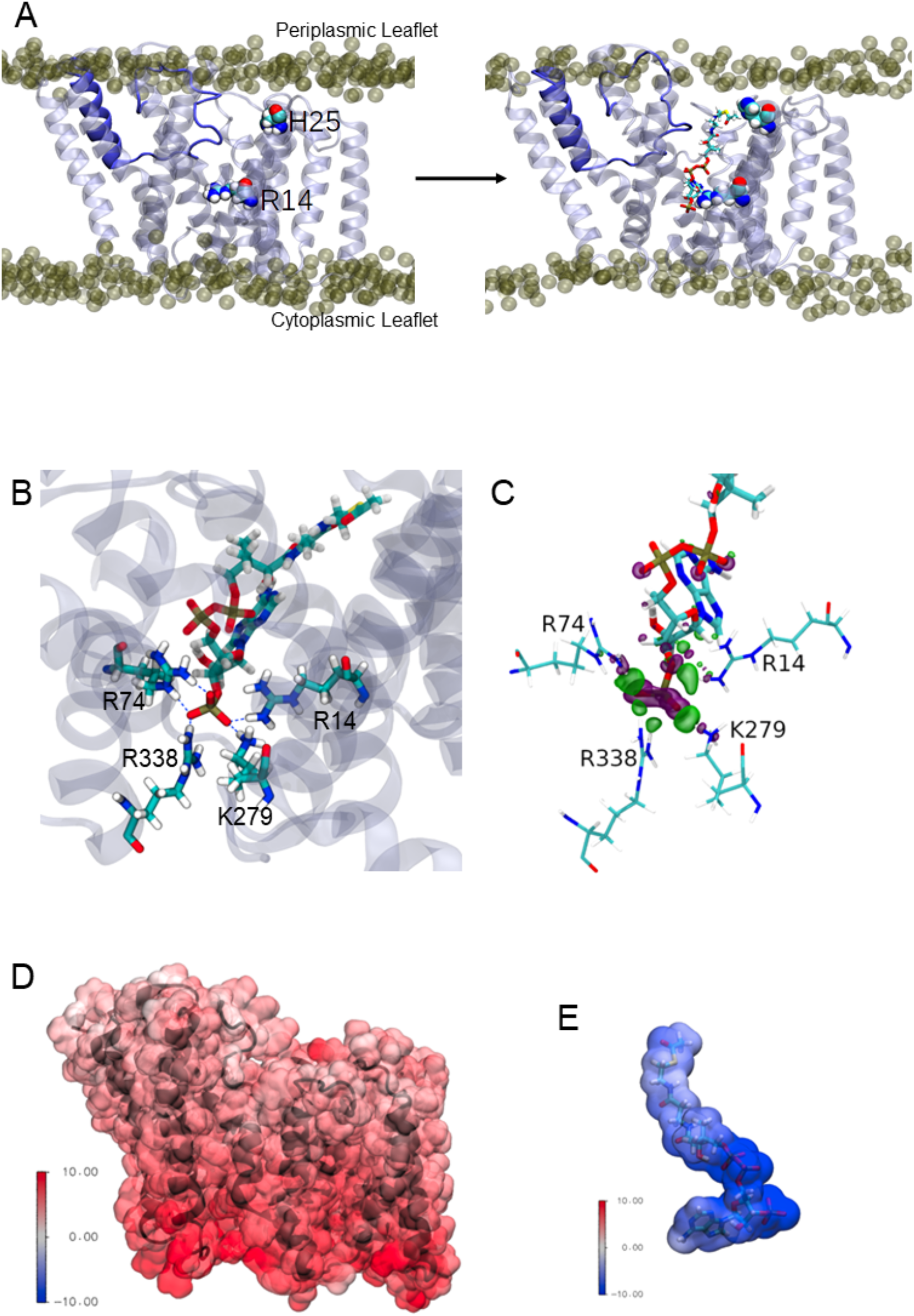
Interactions between the Transmembrane domain of OafB and the putative acetyl donor molecule, acetyl coezyme-A. Note these structures are taken from simulations in which we use the full OafB protein, but only the transmembrane domain (residues 1-377) is shown here for clarity. **(A)** Left: Initial structure of the OafB transmembrane domain. Essential residues H25 and R14 shown as spheres, the loop between helices 3 and 4 is shown in dark blue. Phosphate headgroups of the phospholipids shown as tan spheres. The loop initially occludes the pore in the AT3 domain. Right: Structure from the end of the steered MD simulation in which acetyl Coenzyme-A was pulled into the central channel within the AT3 domain. The loop between helices 3 and 4 has moved away from the center of the pore towards TMH2 (to the left) sufficiently to allow passage of acetyl-coa into the transmembrane domain. **(B)** A pocket of basic residues is observed in the AT3 domain, complementary to the 3’-phosphate of acetyl coenzyme-A. Several high occupancy hydrogen bonds are observed between the 3’-phosphate and transmembrane domain residues R14, R74, R338, and K279. **(C)** Charge transfer interactions between acetyl coenzyme-A and the transmembrane domain of OafB, identified via ONETEP Energy Decomposition Analysis. Loss of electron density is depicted as a purple surface and gain of electron density in a green surface. Significant charge transfer interactions were identified between the 3’-phosphate of acetyl Coenzyme-A (losing electron density) and surrounding basic residues R14, R74, R338, and K279 (gaining electron density) **(D)** Electrostatic potential of the OafB transmembrane domain. Protein in black New Cartoon representation; calculated electrostatic potential overlayed in Surface representation. Scale from -10 V (blue) to +10 V (red). **(E)** Electrostatic potential of acetyl Coenzyme-A. Molecule in Licorice representation; calculated electrostatic potential overlayed in Surface representation. Using the same scale as the AT3 domain, it is clear that acetyl coenzyme-A and the AT3 domain are complementary.

The acetyl-CoA molecule was then allowed to equilibrate within the transmembrane domain using equilibrium MD. Three replicates of this system were generated and simulated with altered starting conformations for the acetyl-CoA within the protein. In the unrestrained final 20 ns of these simulations, acetyl-CoA was observed to hydrogen bond to several basic residues within the pore (**Fig. 5B**). In addition to the Arg14 coordinating the phosphate of 3’-phosphate acetyl-CoA, we noted that the conserved Arg74 (of the RXXR motif of TMH3) along with Lys279 and Arg338 formed a small pocket capable of orienting such that they can all simultaneously hydrogen bond to the 3’-phosphate group (**Fig. 5B**).

Further to this, energy decomposition analysis using ONETEP (86,87), indicates strong, attractive interactions between the AT3 domain and acetyl-CoA. The transmembrane domain-acetyl-CoA complex was optimised, and a single point energy calculated. This energy decomposition analysis approach was employed to calculate the quantum mechanical interaction energy and decompose it into its contributing electronic components accounting for electrostatic, exchange, correlation, Pauli repulsion, polarisation, and charge transfer contributions. Adding the dispersion component, the overall interaction energy was calculated to be -1,035.943 kcal mol-1. Strong charge transfer interactions were identified between the protein and small molecule. The most prominent of these were between the 3’-phosphate of acetyl-CoA and nearby basic residues in the AT3 domain (Arg14, Arg74, Arg338, Lys279); these interactions are shown in **Figure 5C** and are in strong agreement with the above MD simulations.

We calculated the electrostatic potential of the transmembrane domain and acetyl-CoA, and found them to be complementary: the transmembrane domain displayed positive potentials, while the acetyl-CoA displayed negative potentials, of similar magnitudes (**Fig. 5D,E**). The greatest positive potentials are found towards the cytoplasmic side of the transmembrane domain (in agreement with the positive inside rule (88)), and the greatest negative potentials are found at the phosphate groups of acetyl-CoA (which are positioned towards the cytoplasmic side of the AT3 domain). Together these data are strongly supportive of a model whereby acetyl-CoA from the cytoplasmic side is able to penetrate the enzyme to bring the acetyl group into range for use in catalytic transfer to the receptor sugar on the periplasmic face of the membrane.

### Modelling suggests the two domains in OafB could function in a pedal bin mechanism

To learn more about the function of the model OafB proteins, which requires both the AT3 domain and its linked SGNH domain to function, we first constructed a hybrid structure that combined the RaptorX predicted structure of the transmembrane domain (residues 1-376) with the x-ray crystal structure (380-640) of the SGNH-ext (380-421) and the SGNH domain (422-640). Residues 377-379 were added to the N-terminus of the SGNH domain using MODELLER 10.0 (89), and a peptide bond was generated between residues 376 and 377 in ChimeraX (90) with varying C-N-C_α_-C dihedral angles (80, 100, 120, 140°) to generate proteins with differing relative conformations of the AT3 and SGNH domains (henceforth referred to as OafB_80_, OafB_100_, OafB_120_, and OafB_140_). To investigate the flexibility of the hybrid structure we used equilibrium MD simulations (see Methods).

The x-ray crystal structure of the SGNH-ext region of OafB (PDB ID 6SE; residues 380-421) (58) shows it to be structured and interact substantially with the SGNH domain. The biological relevance of this close interaction is supported by the equilibrium MD simulations of OafB: the SGNH-ext remains structured and packed close to the SGNH domain throughout the 4 × 250 ns simulations, indicating this fold is stable under physiological conditions. Unfolding of the SGNH-ext to generate a longer flexible linker - such as that seen in the RaptorX predicted structures of OatA-LM and OatA-SA - is not observed.

A short stretch of only around 10 residues (∼370-380) remains that could form a flexible linker between the transmembrane and periplasmic domains. Indeed, in the MD simulations, the periplasmic domain exhibits a wide range of orientations relative to the transmembrane domain. Principal component analysis (PCA) of the backbone atoms was used to extract the first 2 major motions (accounting for 77% or more of the variance) of the protein in the four equilibrium replicates. The first of these motions was a pedal bin-like action, where the periplasmic domain ‘lid’ opens and closes relative to the transmembrane domain, hinged at the flexible linker. The second motion was the rotation of the SGNH domain within the periplasmic space about the linker. These two motions can be seen in **Videos 1 and 2**.

Despite these motions and the range of relative orientations, the SGNH and AT3 domains were not observed to spontaneously interact in the equilibrium simulations. This does not necessarily mean that the domains do not interact *in vivo*; given the short timescales of MD simulations and the vast conformational space available to the protein, sampling all possible conformations is an intractable problem. To direct computational efforts to the conformational space of interest, elastic network bonds were applied to pull the transmembrane and periplasmic domains together.

Elastic networks are a set of artificial harmonic ‘bonds’ added to a molecular model. These are commonly used in coarse-grained models to maintain the secondary structure of molecules (91); here, we use them to induce and then maintain a change in the tertiary structure of OafB_x_ by pulling the periplasmic and transmembrane domains towards each other. The ‘bonds’ were added between residues likely to be proximal to one another as identified through coevolution analysis (see Methods). Around 75% of co-evolving residues have heavy atoms (carbon, oxygen, nitrogen, sulphur) within 5 Å of one another (92). The side chains of these residues were generally 3-5 Å in length; to avoid artificial re-orientation of the side chains but still bring the residues close enough to interact, elastic bonds of length 10 Å were added between the alpha carbons of the residue pairs. Applying elastic network bonds between the transmembrane and periplasmic domains of the OafB_x_ proteins resulted in the rapid and irreversible ‘closing’ of the protein (the closed state), for the duration of the simulation time that the elastic network bonds were applied.

The elastic network bonds were then removed from these closed state structures, and over the subsequent 100 ns equilibrium MD simulations, all four replicates remained closed throughout. In this configuration, inter-domain interactions, sufficiently stable to maintain the closed state without the elastic network, are identified. Hydrogen bond analysis (distance and angle cutoffs of 3 Å and 20°, respectively) of the subsequent equilibrium simulations revealed a minimum of 35 unique hydrogen bonds between the periplasmic and transmembrane domains across each simulation. When averaged across all four simulations, the number of hydrogen bonds between the two domains at any given time was 3.16 ± 1.67 (**Fig. 6A**). Three of the higher occupancy hydrogen bonds found in all four replicates reside close to the linker region, as shown in **Figure 6B**. Specifically, these are Lys132-Asp387 (all >51% occupancy), Thr386-Asp98 (>25% occupancy), and Ser129-Asp387 (>70% occupancy in 3 of the 4 simulations, 23% in the fourth). Additional hydrogen bonds were observed further from the linker in each system, on the other side of the catalytic triad: for example, Leu408-Glu189 (34% occupancy) and Ser474-Glu189 (21%) are observed in the OafB_100_ system (**Fig. 6C**), but do not form simultaneously (**Supp. Fig. S4**). Taken together, these findings from equilibrium MD simulations identify interactions between the residues in the AT3 membrane domain and the periplasmic domains, supporting the hypothesis that the two domains may cooperate.

**Figure 6:**
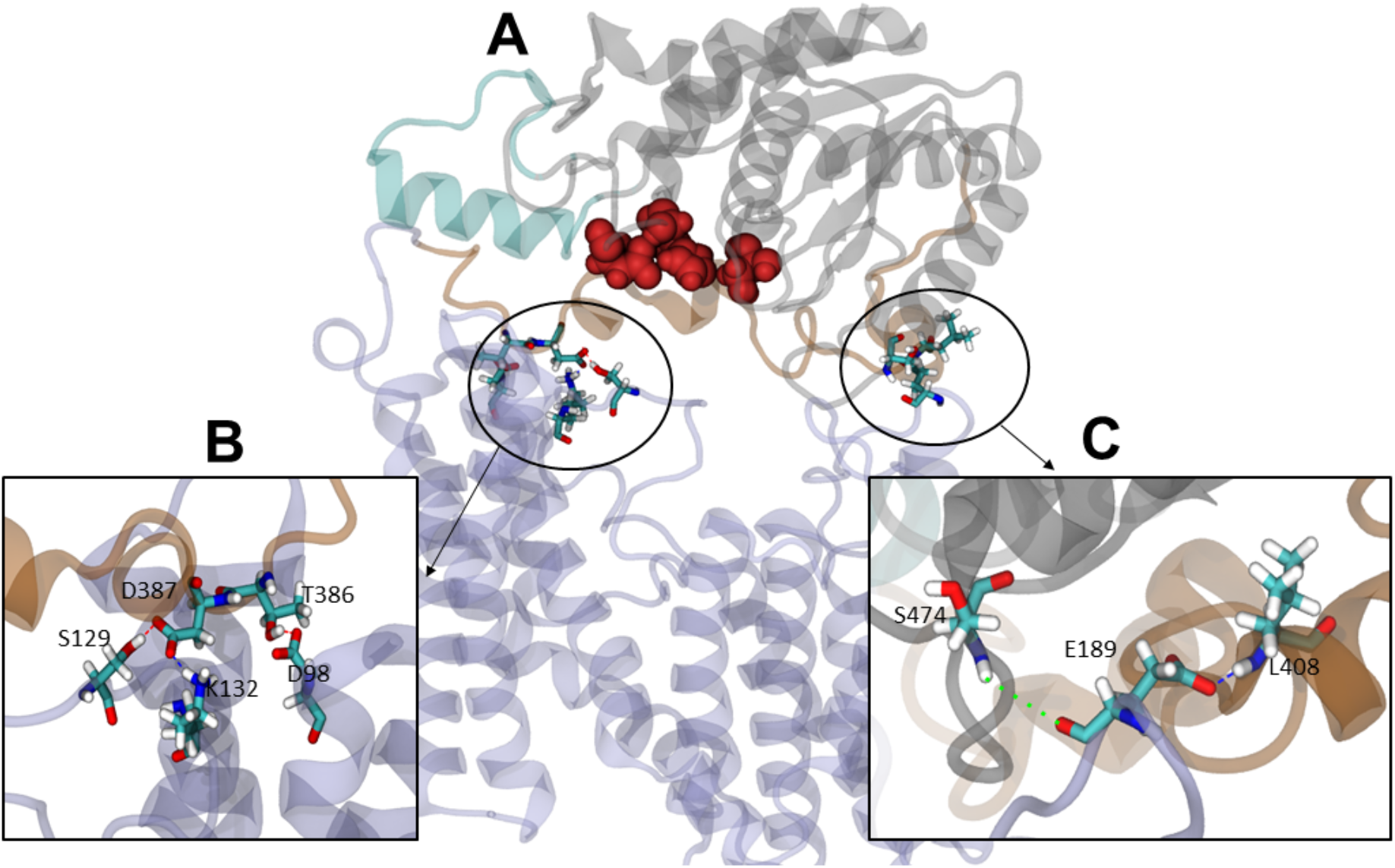
Interactions between the transmembrane and periplasmic domains in the closed state of OafB. **(A)** Closed OafB structure in New Cartoon representation (TM domain in lilac, SGNH-ext in orange, α8 in teal, SGNH domain in grey). Catalytic triad (D618, H621, S430) as red spheres. Residues identified in high-occupancy or important hydrogen bonding interactions between the SGNH and AT3 domains in Licorice representation. **(B)** Hydrogen bonding observed between SGNH-ext and AT3 domains in all equilibrium simulations of the closed OafB structure. High-occupancy hydrogen bonding between side chain: carboxylate of E387 and hydroxyl of S129, amine of K132; carboxylate of E98 and hydroxyl of T386. **(C)** Hydrogen bonding between the SGNH and AT3 domains observed in the OafB100 simulations. E189 can hydrogen bond to S474 via its backbone carbonyl (green dashed line), or to L408 via its carboxylate side chain (blue dashed line) but cannot form these interactions simultaneously. Timeseries of the separation of these residues shown in Supplementary Figure S4.

These fully closed structures were submitted to the ProPores2 webserver (93) to identify and parameterise cavities within the protein. The closing of the pedal bin converts the outer cavity from the transmembrane domain into a larger enclosed cavity at the interface of the AT3 and SNGH domains (**Figure 7A**) This pore has a volume of ∼4,780 Å^3^ and includes both the SGNH catalytic triad comprising residues Asp618, His621 and Ser430, and at the membrane surface the proposed catalytic residues of the AT3 domain, namely, His25 and Tyr194. This raises the possibility that the transfer of the acyl-group to the acceptor molecule could occur in this region of the protein.

**Figure 7:**
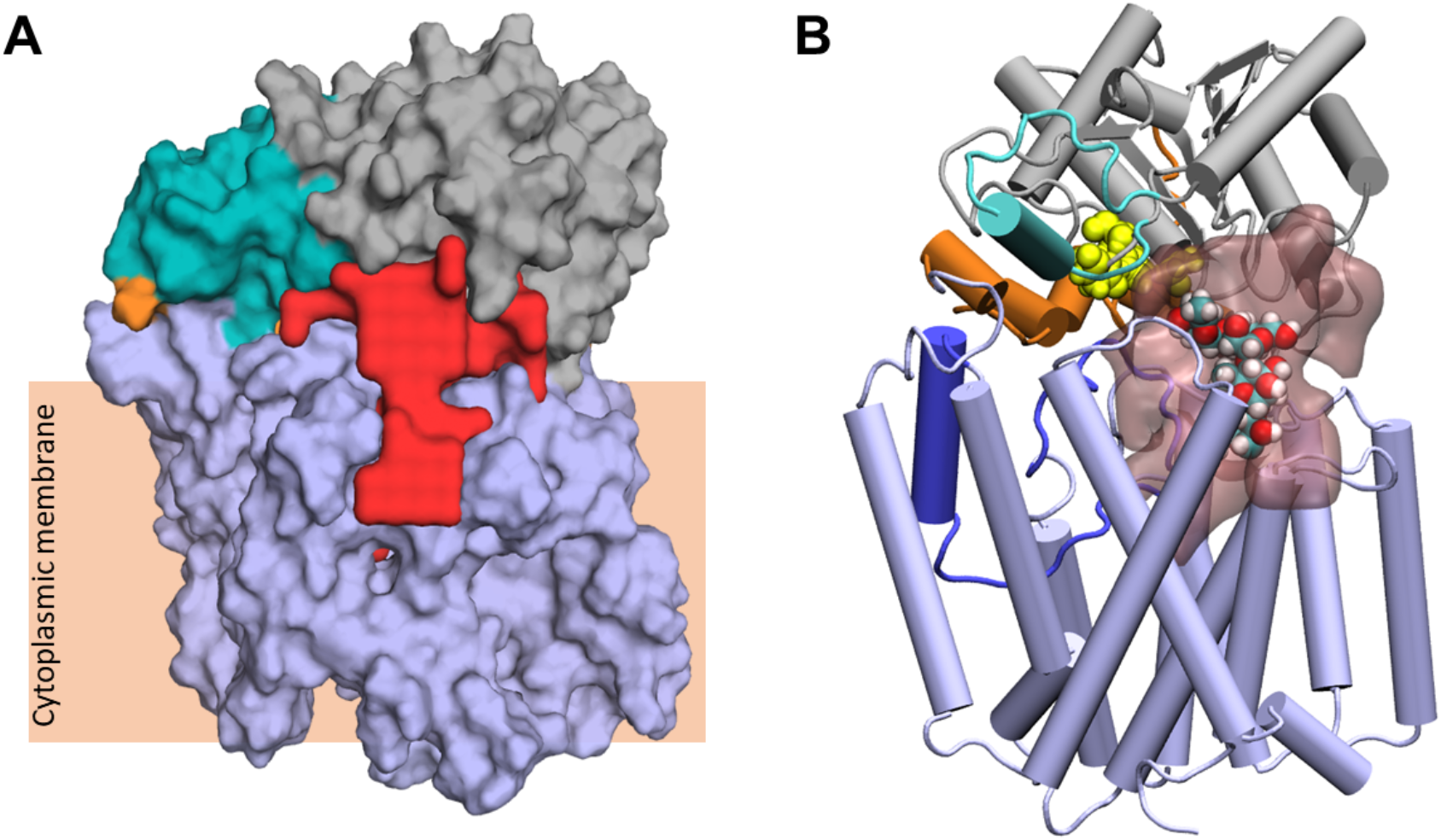
Largest cavity in the closed state of OafB. **(A)** Largest cavity identified by the ProPores2 webserver in the closed OafB structure shown as a red surface. The cavity extends down into the transmembrane domain. **(B)**. Closed OafB in Cartoon representation: transmembrane domain coloured light blue, periplasmic loop between TMH 3-4 coloured dark blue, SGNH domain coloured grey with the periplasmic linking region in orange, and additional helix in the SGNH domain in teal. The SGNH catalytic triad (Ser430, Asp618, His621) is shown as yellow spheres. Largest pore identified by the ProPores2 Webserver shown as a transparent pink surface. A single O-antigen unit is shown inside the pore in van der Waals representation, coloured by atom name (cyan for carbon; red for oxygen; white for hydrogen). The single O-antigen unit can be comfortably accommodated in this space in several orientations.

### Fully closed structure exhibits cavity able to accommodate the LPS O-antigen

Finally, we considered the potential mechanisms by which the acceptor molecule(s) would interact with open and closed ‘pedal bin’ model of OafB, working with two alternative hypotheses that the acceptor is presented either as a single or multiple O-antigen repeat unit(s) anchored to a lipid carrier or later after transfer to the lipidA core molecule.

The closed form of the pedal bin model presents a possible enclosed catalytic site for OafB, which is known to acetylate the rhamnose residue in the O-antigen repeating unit of the LPS (10). This cavity resides directly below the catalytic triad (Ser430, His621, Asp618), and extends down into the AT3 domain (**Fig. 7A**). The measured volume of ∼4,780 Å^3^ is an order of magnitude greater than the volume of the LPS O-antigen unit (volume of ∼502 Å^3^), hence one or more units could be manually placed within the cavity (**Fig. 7B**). Importantly the enclosed cavity on the periplasmic surface of the membrane does extend into the membrane, the lower portion residing below the plane of the upper leaflet headgroups and hence is exposed to the membrane core (**Fig. 7A**), consistent with a ligand delivered on a lipid anchor.

*S. enterica* serovars Typhimurium and Paratyphi use the prevalent Wzx/Wzy-dependent pathway for O-antigen assembly, in which a single O-antigen unit is assembled in the cytoplasm on undecaprenyl phosphate (Und-PP) in the membrane, which is then flipped placing the O-antigen unit in the periplasm. Lipid linked O-antigen units are assembled to the final length in the periplasm before ligation to lipid A-core. Subsequently, the completed LPS molecule is transported to the outer membrane (94). Thus, the question remains, does OafB acetylate the rhamnose moiety of a single O-antigen unit bound to Und-PP or after O-antigen polymerisation? If the latter, does acetylation occur with the polymerised OAg unit(s) attached to the Und-PP or after transfer to the lipidA-core molecule?

To contextualise the OafB models with biologically relevant molecules, a system containing one molecule of the lipid carrier undecaprenyl phosphate (Und-PP) attached to an O-antigen unit was placed next to the OafB model (**Fig 8A**). A distance of ∼9 Å separates O3 of the rhamnose from the periplasmic surface of the membrane, which is similar to the distance of the SGNH active site (Ser430) from the membrane (which was ∼12 Å calculated with OafB_100_ after 100 ns with elastic network bonds applied), meaning it is possible that the substrate diffuses laterally into the cavity, either with the pedal bin open or closed.

**Figure 8:**
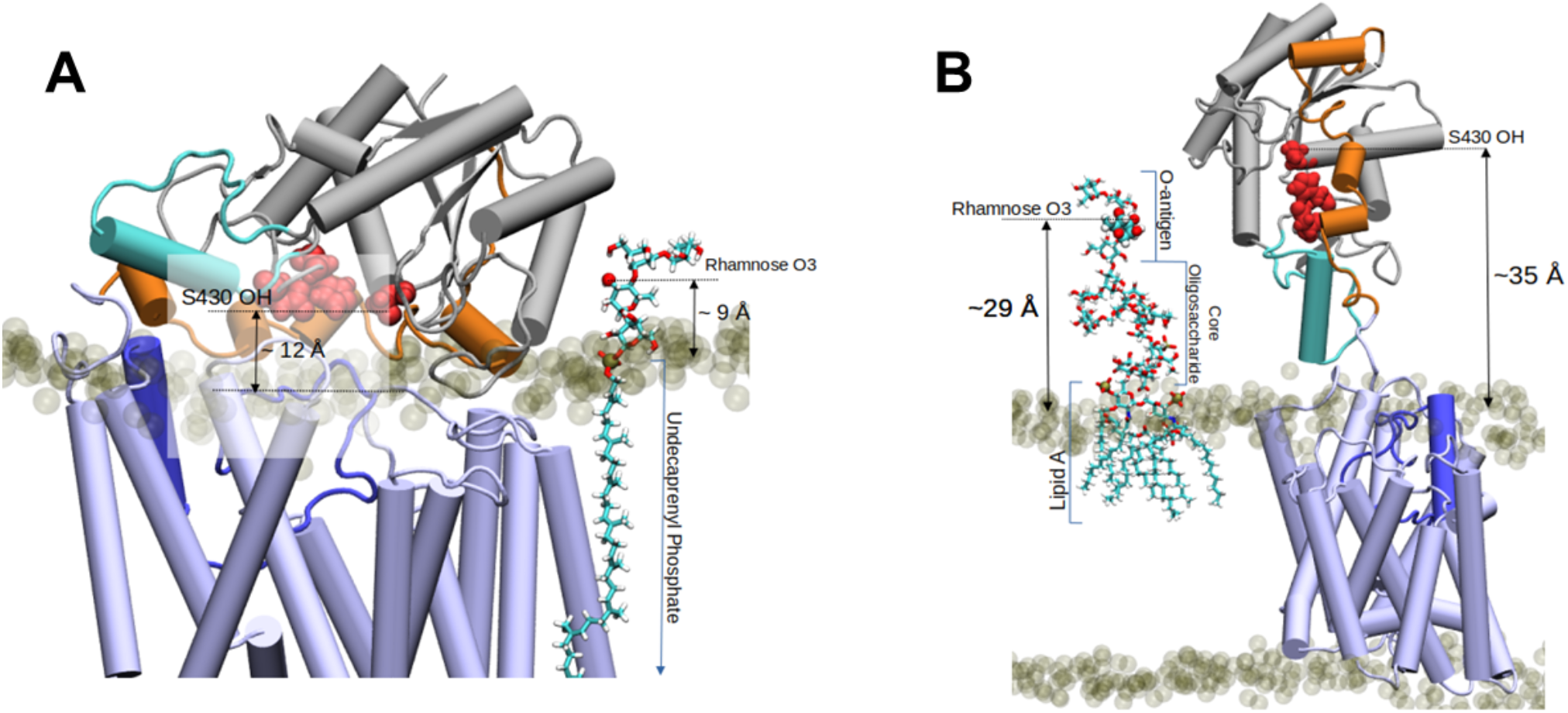
Two proposed models for the O-antigen rhamnose moiety acetylation by OafB. In both panels, OafB is shown in a cartoon representation with the AT3 domain coloured light blue, periplasmic loop between TMH3-4 coloured blue, SGNH domain coloured grey, the periplasmic linking region in orange, and additional helix in the SGNH domain in teal. Catalytic triad of the SGNH domain (Ser430, Asp618, His621) shown as red spheres. POPE and POPG phosphate headgroups shown as transparent tan spheres. **(A)** Undecaprenyl phosphate carrier lipid with a single O-antigen unit in Licorice representation. O3 of the rhamnose moiety (marked by a red sphere) has a vertical separation from the periplasmic leaflet surface of ∼9 Å. The OH of S430 has a vertical separation from the bulk membrane of ∼12 Å. **(B)** OafB-LPS system after equilibration steps - one LPS molecule with a single O-antigen unit shown in Licorice representation, and its rhamnose moiety shown in van der Waals representation. OafB remains in an open conformation. After relaxation, the oligosaccharide of LPS has kinked, with the distance between the lipid headgroups of the periplasmic leaflet surface and the acetylation site reduced to ∼29 Å (from ∼42 Å, as shown in Figure S5). This is comparable to the ∼35 Å separation between the OH of S430 and the periplasmic leaflet surface

The analysis above suggests that acetylation could occur using single O-antigen units, before O-antigen polymerisation. Alternatively, acetylation occurs during or after polymerisation on the Und-PP carrier before transfer to the lipidA-core (not shown). The distance of multimers of 9 Å (single O-antigen unit) stretching away from the membrane could possibly be accommodated by the open state of the pedal bin with the SGNH domain extending into the periplasm.

To investigate the final model of acetylation after assembly onto the lipidA core, we built a model with LPS (with one O-antigen unit) and OafB-SPA embedded in the inner membrane to assess the distances defined between the periplasmic leaflet surface and various groups within the system relevant to the acetylation (**Fig. 8B**). One molecule of *Salmonella* LPS was generated in CHARMM-GUI’s LPS Modeler and inserted into the periplasmic leaflet of the OafB_100_ system (replacing 2 POPE and 2 POPG molecules). Initially the modelled LPS adopts a quasi-linear conformation; the z-separation between the rhamnose moiety and the periplasmic membrane surface was found to be ∼42 Å, as shown in **Supplementary Figure S5**. During minimisation and equilibration, the oligosaccharide was allowed to relax; this resulted in a kinked structure, where the distance between the acetylation site and the periplasmic leaflet surface was reduced to around 29 Å (**Fig. 8B**). By comparison, after equilibration, the z-separation of the hydroxyl of Ser430 in the catalytic triad of OafB_100_ and the periplasmic leaflet is ∼35 Å, hence it is not inconceivable that the open pedal bin could function to deliver acetyl groups at a distance from the membrane. However, we note that in reality the full-length O-antigen will be attached to the lipidA core, which would require significant, entropically unfavourable conformational rearrangement of the O-antigen for OafB to catalyse significant overall levels of O-acetylation across the whole polymer.

Steered molecular dynamics (SMD) was used to pull the rhamnose moiety of LPS and Ser 430 of OafB towards one another, and subsequent restrained MD simulations held the protein and LPS close to one another to assess possible interactions. Many hydrogen bonds were observed between the O-antigen and the periplasmic domain of OafB over the two replicates. Several of these were with residues from the SGNH-ext (galactose-Asp387; galactose-Tyr389; rhamnose-Tyr394). Tyr389 and Tyr394 are both residues which were previously identified by FTMap *in silico* docking analysis as involved in binding to rhamnose sugars. Hydrogen bonding was also observed between the 3’ and 2’ hydroxy groups of the rhamnose moiety and Ser430. Even when all position restraints were removed in the MD simulations, this interaction persisted for a further 10 ns.

Together these analyses indicate that O-acetylation of the O-antigen may occur either when a single O-antigen unit attached to a lipid carrier is presented in the periplasm, or when the O-antigen is polymerized, either on the lipid carrier, or when attached to the lipid A-core.

### Ideas and Speculation

The realisation that diverse bacteria use AT3 proteins in a plethora of different biological processes for the modification of extracytoplasmically localised glycans, directed our efforts into describing the fundamental mechanism of these proteins (1). While the role of AT3 proteins in these diverse processes is often implicated from genetics only, the OafB protein has been studied in much more detail, along with other selected AT3 proteins including the OatA protein from *S. aureus* (10,57,58). These systems are examples of AT3 proteins working with SGNH domains to acylate LPS and peptidoglycan, respectively, which are key components of bacterial envelopes.

Integration of experimental and computational data has provided detailed insight into the structure and potential mechanism of action of AT3-SGNH proteins. Our structural model, combined with molecular dynamics simulations, suggests first that the transmembrane domain of OafB has a novel fold. Importantly this is distinct from the MBOAT family of membrane-bound acyltransferases, suggesting that nature has evolved two independent routes to solve this problem. The discovery of a large cytoplasmic cavity, which can open dynamically to accommodate an acyl-CoA molecule that spans the membrane, provides a key mechanistic breakthrough in our understanding of the protein function, which in OafB is mediated by a loop region which would need to block the pore in the absence of donor/acceptor to prevent leakage of protons.

The putative catalytic residues His25 and Tyr194 are close enough to the acetyl-group (∼ 5 Å) to conceivably be involved in catalysis, although the MD simulations cannot capture bond making or breaking (nor proton transfer). The loop between TMH 5 and 6, in which Tyr194 resides, shows significant mobility and the side chain of Tyr194 is observed to point inwards, towards the centre of the pore (and therefore towards acetyl-CoA). Given the conformational mobility of the 4’-phosphopantetheine segment of the coenzyme, and of the loop between TMH 5 and 6, it is not unreasonable to suggest that the thioester bond is accessible to both His25 and Tyr194 to aid the acetyl transfer in a mechanism such as that suggested by Jones et al (57). Addressing the role of Glu334, which Jones et al (56) proposed was the third residue in the catalytic site (Glu357 in OatA), our model places this distantly from the His25 and Tyr194, but yet in OafB we also know this residue is important for function (79). The residue sits close to the cytoplasmic side and the internal cavity and perhaps has a role in the initial recognition of the acyl-CoA. Other conserved residues, including Phe42, Tyr124, Trp138 and Glu143 in OafB, which were identified by mutagenesis as being important for both the OafB-related protein OafA and OatA, cluster on the extracytoplasmic surface and could have related roles in acyl-CoA binding, catalysis or acceptor recognition.

While the function of the AT3 to bind and deliver acyl-groups to a periplasmic catalytic site is well supported by our model, and must be a conserved feature of all AT3 proteins, the further discrete role of the SGNH domain is still not fully understood, although is essential for transferase function in both OafB and OatA (57,58). The linker region, while short and structured in OafB suggests that the domain can function in a ‘pedal bin’ like model, opening and closing to define a large cavity on the extracytoplasmic surface, which we demonstrate could then capture a lipid-anchored O-antigen repeat structure. For OatA and other related proteins, this linker can be much longer (**Fig. 2**) and whether this then allows the pedal bin lid to close and accept the acyl-group and then open and extend away from the membrane to reach the acceptor substrate needs further investigation, but the MD simulations suggest it is possible. However, for the OafB system we favour the model of lid opening and closing to help trap the substrate in a membrane-surface-located activity site, similar to the function of the Sus protein in capturing ligand in the outer membrane of *Bacteroidetes* bacteria (95). For LPS O-acetylation by OafB and similar proteins we favour the model where the substrate is the single O-antigen repeat bound by the Und-PP lipid anchor before later polymerisation. Given that overall levels of OafB-dependent O-acetylation vary between 40-70% (10,96), the stochastic interaction of OafB with the Und-PP anchored O-antigen repeat appears a more parsimonious mechanism than relying on the enzyme trying to reach rhamnose sugars in a fully assembled O-antigen.

Beyond LPS and peptidoglycan, AT3 domain-containing proteins acylate a diverse range of complex carbohydrates in bacteria (1). Each receptor substrate may present its own unique challenges for the AT3 domain. In both the LPS and peptidoglycan examples, additional SGNH domains are required for function, however, AT3 domains can function as standalone proteins. One example of this is IcaC, which is involved in the O-succinylation of poly N-acetylglucosamine (PNAG) biofilm component of *Staphylococci*. Here, the acceptor is not delivered on a lipid anchor, but rather through a continuous synthesis and extrusion mechanism catalysed by IcaAD, a ‘synthase’ mechanism (94). Perhaps a close physical association of IcaC with IcaAD removes the need for the additional SGNH domain and allows direct transfer of the acyl group onto the emerging PNAG polymer giving stochastic levels of O-succinylation of about 40% (97,98). Hence, the differences between AT3 and AT3 with additional domains such as SGNH, perhaps could relate to the route of presentation of the acceptor molecule within the context of different biosynthetic pathways for extracytoplasmic glycans.

## Conclusions

Here we provide strong evidence that the integral membrane AT3 domain (PF01757), mediating acylation of a diverse range of complex carbohydrates, has a novel fold. The stability of this structure was confirmed using molecular dynamics simulations, which also enabled insights into the mechanism of action of AT3 domains. Importantly, for the first time our model provides a solution to the problem of acetylation occurring in the periplasm using the cytoplasmic acetyl donor acetyl-CoA. We show a membrane-spanning pore occurs transiently in the AT3 domain which can accommodate acetyl-CoA, presenting the acyl group to the periplasmic side. This part of our model is supported by quantum-level calculations to probe the hypothesised protein-acetyl-CoA interactions, and the presence of key, conserved and essential residues that line the pore. The second cavity at the periplasmic side in OafB is able to accommodate the receptor substrate O-antigen. Together, our data give important new insights into the family of AT3 domain containing proteins. Our well-supported model can support and drive targeted research to gain a full understanding of the mechanism of action of this family or membrane bound acyltransferase proteins that mediate acylation of complex carbohydrates across the domains of life.

## Methods

### Co-evolution and protein structure predictions

OafB Protein sequence (Uniprot accession A0A0H2WM30) was submitted to either the RaptorX contact prediction server (66–68) or AlphaFold server (76) and analysis was run with default parameters. Predicted structures using RCSB PDB Pairwise Structure Alignment webserver, employing the flexible jFATCAT alignment algorithm (99).

### Molecular dynamics simulations

All simulations used the GROMACS simulation package (version 2020.3) (100). Results were analysed using GROMACS tools and in-house python scripts utilising MDAnalysis (101–104). Molecular graphics were generated using VMD 1.9.4 (105).

### Atomistic simulations

Atomistic simulations used the CHARMM36m forcefield (106) with the TIP3P water model (107). A cut-off of 1.2 nm was applied to Lennard-Jones interactions and short-range electrostatics using the potential shift Verlet scheme. Long-range electrostatics were treated using the particle mesh-Ewald (PME) method (108). Atoms were constrained using the LINCS algorithm to allow the use of a 2 fs timestep (109). For production simulations, temperatures were maintained using the Nosé-Hoover thermostat (110,111) (1.0 ps coupling constant), and pressure was maintained at 1 bar using the Parrinello-Rahman semi-isotropic barostat (112) (5.0 ps coupling constant).

### Transmembrane Domain Model

A model of the transmembrane domain (residues 1-376, i.e. the 10 TMH of the AT3 domain and the 11th TMH) was generated using the RaptorX webserver (66,67). Using the CHARMM-GUI membrane builder module (113), this structure was embedded within a model *Escherichia coli* inner membrane: a symmetric bilayer of 18:1:1 1-palmitoyl-2-oleoyl-sn-glycero-3-phosphoethanolamine (POPE), 1-Palmitoyl-2-oleoyl-sn-glycero-3-phosphoglycerol (POPG), and 1’,3’-bis[1-palmitoyl-2-oleoyl-sn-glycero-3-phospho]glycerol (cardiolipin). The bilayer system was subsequently solvated in 0.15 M KCl. The system was energy minimised in 5000 steps using the steepest descent method (114). The subsequent structure was equilibrated in six phases in which the protein and lipid head groups were subjected to position restraints with varying force constants. The full equilibration protocol is detailed below. Two short, independent equilibrium molecular dynamics simulations were carried out on the equilibrated system: one simulation of 100 ns at 303.15 K, and one of 50 ns at 320 K.

### Equilibration Protocol

Each system was equilibrated using two NVT phases followed by four NPT phases. NVT phases used a timestep of 1 fs and each lasted 0.125 ns, and the NPT stages used a timestep of 2 fs and each lasted 0.5 ns. The velocity-rescaling thermostat (115) was applied at all stages to bring the system to 303.15 K with a coupling constant of 1.0 ps. Semiisotropic Berendsen pressure coupling (116) was applied to the NPT phases to equilibrate with a pressure bath of 1 bar (τ_P_ = 5.0 ps, compressibility of 4.5×10^−5^ bar^-1^).

### Full OafB model

The RaptorX transmembrane domain model and periplasmic domain crystal structure (PDB ID: 6SE1) were used to build a full OafB model. Missing linker residues (residues 377-379) were modelled into the N-terminus of the SGNH domain using MODELLER 10.0 (89). A peptide bond was generated between the N-terminus of the resulting periplasmic domain (K377) and the C-terminus of the transmembrane domain (N376) using UCSF ChimeraX (90). Four structures were generated, varying the C-N-C_α_-C dihedral angle; values of 80, 100, 120, and 140° were used to give structures with different relative orientations of the two domains. The C_α_-C-N-C_α_ dihedral was maintained at 170°. These proteins will henceforth be referred to as OafB_x_, where x is the C-N-C_α_-C dihedral angle (80, 100, 120, or 140°).

The transmembrane domain of each model was embedded in an *Escherichia coli* inner membrane model and solvated in the CHARMM-GUI membrane builder as described for the RaptorX model. Systems were energy minimised in 5000 steps using steepest descent algorithm, and subsequently equilibrated using the protocol described above. Duplicates of each equilibrated system were generated. The first replicate of each system was simulated for 250 ns at 303.15 K. The topologies of the second replicates were modified to add elastic network bonds that would bring the periplasmic and transmembrane domains together. Elastic bands (of length 1 nm and strength 100 kJ mol^-1^ nm^-2^) were generated between the alpha carbons of 10 residue pairs, identified through coevolution analysis, to be more than 50% likely to be proximal to one another.

These four replicates were simulated for 100 ns with the elastic network bonds applied. A subsequent 100 ns simulation was undertaken on each system with the network removed.

### Full OafB Model: AlphaFold

For comparison to the hybrid models built above, the OafB amino acid sequence was submitted to AlphaFold (76) to generate a structure prediction for the full protein. The RCSB PDB Pairwise Structure Alignment web service (99) was used to compare our structures to the AlphaFold prediction. Residues 1-327 of the RaptorX model, residues 328-406 of OafB80, and residues 407-640 of the periplasmic domain crystal structure (PDB ID: 6SE1) were aligned to the equivalent residues in the AlphaFold model using the flexible jFATCAT algorithm.

### Acetyl Coenzyme A

One molecule of acetyl coenzyme A (acetyl-coA) was generated in CHARMM-GUI’s Ligand Reader & Modeler (117). Using VMD as a visual tool to guide positioning, the acetyl group of acetyl-coA was placed beneath the gap between transmembrane helices 9 and 10, and the N-terminus of the OafB_100_ protein. The protein was embedded in the *Escherichia coli* inner membrane model in CHARMM-GUI and solvated in 0.15 M KCl. The system was energy minimised using the steepest descent algorithm, and subsequently equilibrated using the protocol described above.

A short steered molecular dynamics (SMD) simulation was used to insert acetyl-coA into the pore (delimited by transmembrane helices 1, 2, and 9, and the periplasmic loops between helices 5 and 6, and 3 and 4). Pulling at a rate of 0.5 nm ns^-1^ and with a harmonic force of 1000 kJ mol^-1^ nm^-2^, the sulphur of the acetyl-coA molecule was pulled upwards, towards the centre of mass of the alpha carbons of residues E243, D126, E189 (all in loops at the periplasmic surface of the transmembrane domain, surrounding the pore) until the z-separation of the two groups was 0 (7.5 ns).

Three variations on the final frame of the SMD were generated in VMD by rotating and translating acetyl-coA within this pore to use as starting conformations for equilibrium MD. These systems were energy minimised using the steepest descent algorithm to resolve steric clashes, and equilibrated as described above, with additional position restraints (**Table 3**) on the acetyl-coA molecule. Each system was subjected to three consecutive MD simulations of 20 ns at 303.15 K, with decreasing position restraint strength on acetyl-coA to allow the protein to relax around the substrate.

**Table 1:**
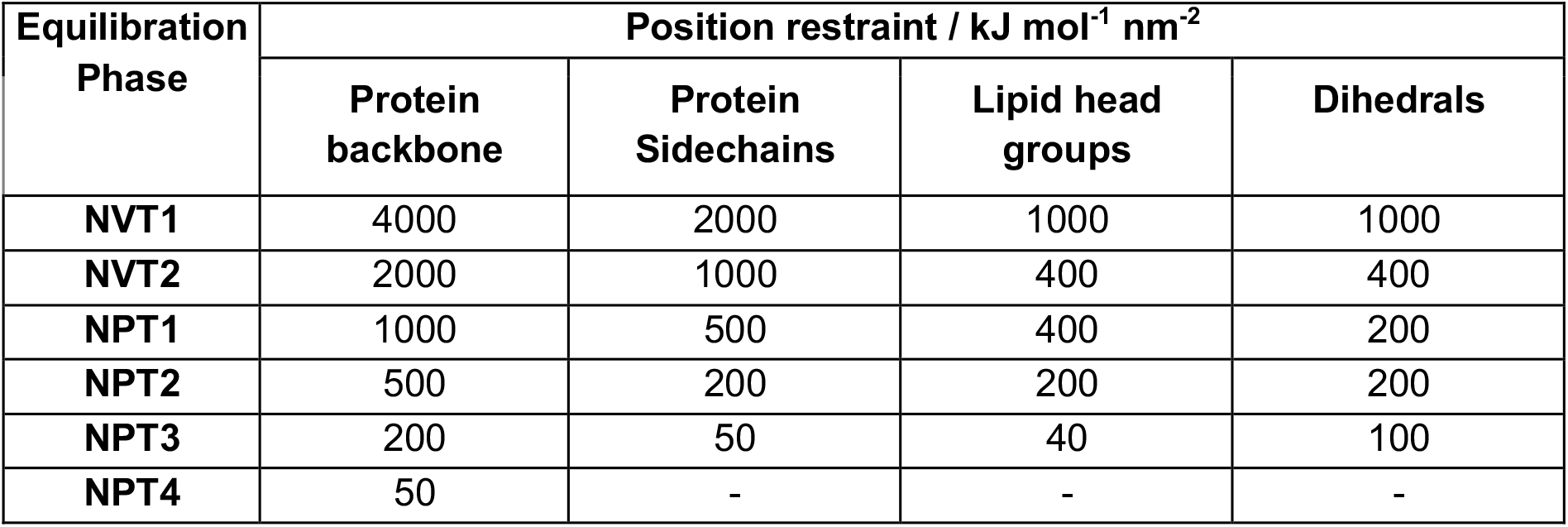
Position restraints used in the equilibration steps for all atomistic systems

**Table 2:** Residues identified through co-evolution as likely to be proximal to one another

| Residue 1 (TM domain) | Residue 2 (periplasmic domain) | % Probability |
| --- | --- | --- |
| 94 | 545 | 56.1 |
| 95 | 546 | 80.5 |
| 96 | 546 | 57.2 |
| 97 | 546 | 70.5 |
| 98 | 546 | 58.4 |
| 123 | 458 | 53.3 |
| 125 | 403 | 59.2 |
| 126 | 404 | 55.9 |
| 194 | 627 | 50.9 |
| 242 | 503 | 51.3 |

**Table 3:** Position restraints used in the equilibration and simulation of the OafB-acetyl CoA systems.

| Equilibration Phase | Position restraint / kJ mol<sup>-1</sup> nm<sup>-2</sup> |  |  |  |  |
| --- | --- | --- | --- | --- | --- |
|  | Protein backbone | Protein Sidechains | Lipid head groups | Dihedrals | Acetyl-coA |
| **NVT1** | 4000 | 2000 | 1000 | 1000 | 4000 |
| **NVT2** | 2000 | 1000 | 400 | 400 | 4000 |
| **NPT1** | 1000 | 500 | 400 | 200 | 4000 |
| **NPT2** | 500 | 200 | 200 | 200 | 3000 |
| **NPT3** | 200 | 50 | 40 | 100 | 3000 |
| **NPT4** | 50 | - | - | - | 2000 |
| **MD1** | - | - | - | - | 1000 |
| **MD2** | - | - | - | - | 500 |
| **MD3** | - | - | - | - | - |

### LPS in the Periplasmic Leaflet of the Inner Membrane

One molecule of *Salmonella spp*. LPS was generated using the CHARMM-GUI LPS modeller (O9-4 core with one O-antigen unit). Using VMD as a tool to guide positioning, 2 POPE and 2 POPG molecules were removed from the periplasmic leaflet of the OafB_100_ system and the LPS lipid A moiety inserted in their place. 8 water molecules were replaced by potassium ions to neutralise the additional negative charge of the LPS. The system was energy minimised in 5000 steps via the steepest descent algorithm to resolve steric clashes. The system was equilibrated in 6 phases, as described for the other atomistic systems.

Steered MD was used to guide the rhamnose moiety of the LPS towards the Serine 430 residue (2 replicates, one with a pull rate of 0.5 nm ns^-1^, the other with a pull rate of 1 nm ns^-1^; both with a pull force of 500 kJ mol^-1^). The final frame of each of these trajectories (13.1 ns and 6.5 ns, respectively) was used as the starting conformation for 3 consecutive MD simulations at 303 K of 20 ns each, with varying position restraints on the protein backbone (250, 0, 0 kJ mol^-1^) and LPS molecule (1000, 500, 0 kJ mol^-1^). Hydrogen bond analysis was undertaken in VMD (distance cutoff = 3 Å, angle cutoff = 20°) to assess the interactions between the LPS and the SGNH domain.

### DFT Calculations

The electrostatic potentials of the transmembrane domain of OafB (residues 1-370) and the acetyl-CoA molecule were calculated in a vacuum using the linear-scaling density functional theory package, ONETEP (86,87), using the PBE exchange-correlation functional (118), augmented with Grimme’s D2 dispersion correction (119). Open boundary conditions *via* real-space solution of the electrostatics were used in a simulation cell with dimensions 9 nm × 7 nm × 8 nm. Norm-conserving pseudopotentials were used for the core electrons, and the psinc basis set, equivalent to a plane wave basis set with a kinetic energy cut-off of 800 eV, was employed. 8.0 Bohr localisation radii were used for the nonorthogonal generalised Wannier functions (NGWFs). The energy decomposition analysis (EDA) implemented in ONETEP (120,121) was used to calculate the intermolecular interaction energy between Acetyl-CoA and the AT3 domain and to extract its contributing components and to visualise charge transfer interactions.

## Supporting information

Video 1 OafB motion

Video 2 OafB motion

## Data availability

The trajectories generated and the run input files necessary to repeat the simulations presented here are openly available in Zenodo at https://doi.org/10.5281/zenodo.6834637.

## Acknowledgements

The authors acknowledge access to the following High Performance Computing resources: Iridis 5 at the University of Southampton and the JADE Tier 2 facility (EPSRC grant no. EP/T022205/1) to which access was granted via HECBioSim, the UK High-End Computing Consortium for Biomolecular Simulation (EPSRC grant no. EP/R029407/1); the C. Skylaris group at the University of Southampton for help with ONETEP.

K.E.N. was supported by a Ph.D. Studentship from the Engineering and Physical Sciences Research Council (Project Number: 2446840); S.N.T. was supported by a Ph.D. studentship from the Biotechnology and Biological Sciences Research Council White Rose Doctoral Training Program (BB/M011151/1), “Mechanistic Biology and its Strategic Application.” SLM acknowledges the support of the Federation of European Biochemical Societies (FEBS) through a long-term fellowship.

