## Supplementary Figures for "A novel fold for acyltransferase-3 (AT3) proteins provides a framework for transmembrane acyl-group transfer"

### Supplementary Figures S1-5

#### Supplementary Figure S1

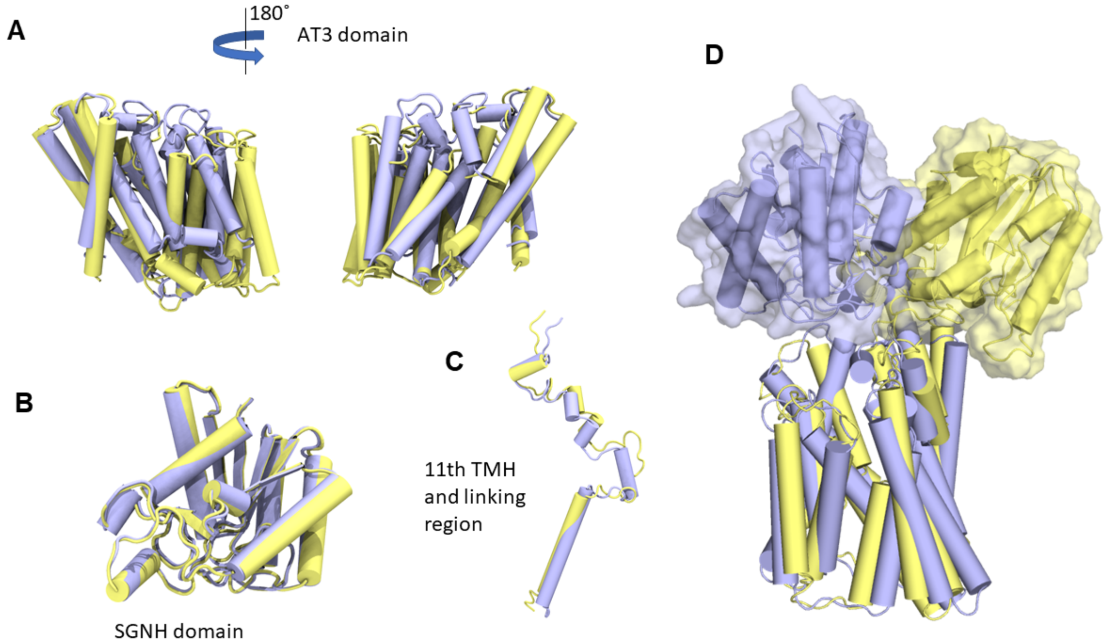


OafB structure as predicted by RaptorX (yellow) and AlphaFold (lilac). Domains (according to the topology in Figure 1A of ref (58)) were isolated from each model and directly compared using a least-squares fit: **(A)** AT3 domain (Pfam PF01757) (residues 1-338); **(B)** SGNH domain (residues 422-640); **(C)** 11th TMH and the periplasmic linking region (residues 328-421). All three domains are largely similar across the two models. **(D)** The RaptorX and AlphaFold models were superimposed. Variation in linker orientation between the two models results in the SGNH domains further apart than one might expect. This supports the idea of a flexible, mobile linker between domains.

#### Supplementary Figure S2

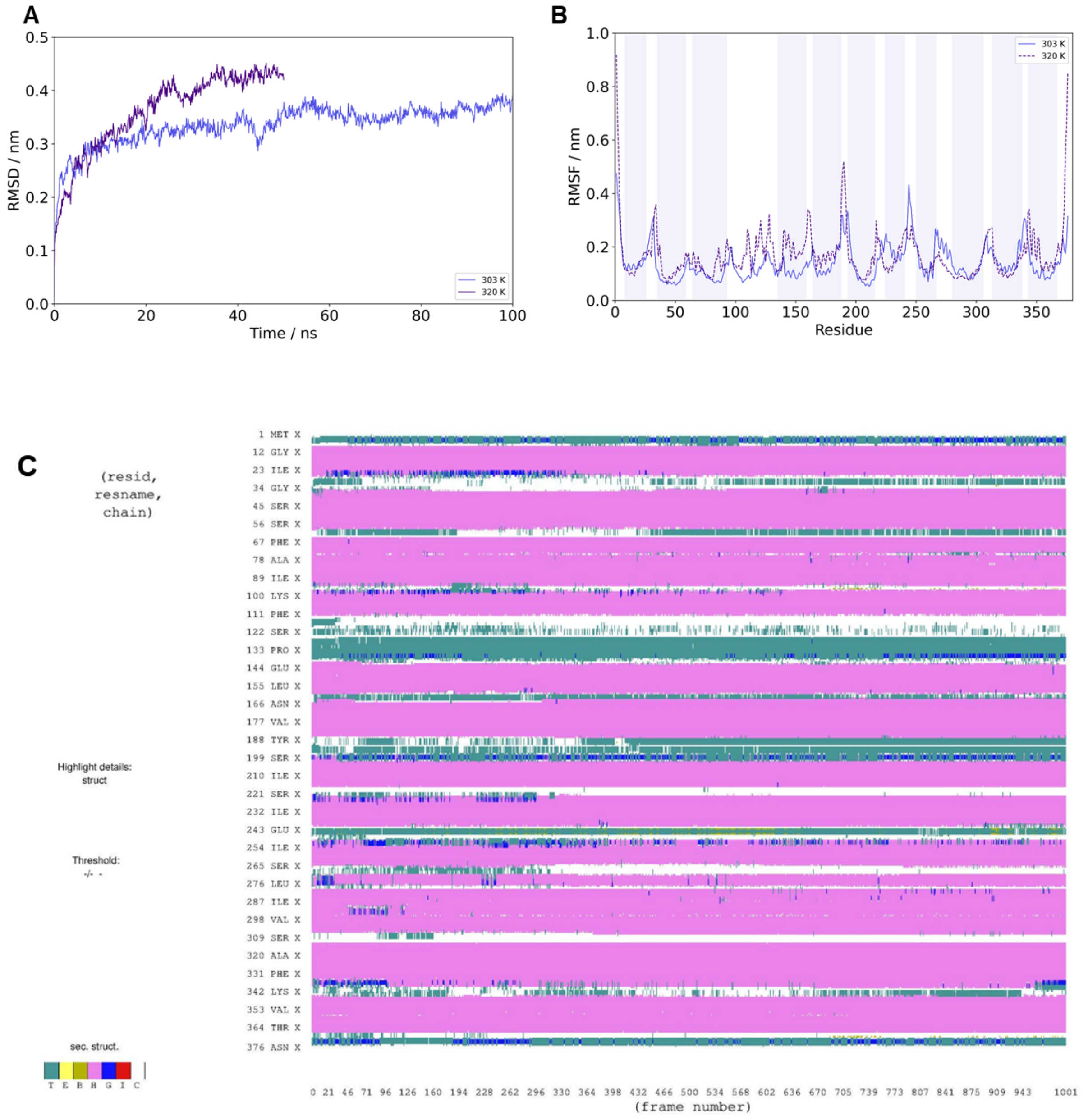


Analysis of the RaptorX model for the transmembrane domain of OafB, embedded in a model *E. coli* membrane and simulated under equilibrium conditions. **(A)** Root-mean-square deviation (RMSD) of the backbone atoms. RMSD values remain below 1 nm for the duration of each simulation, even at an elevated temperature (320 K), indicating that there are no major structural changes occurring. **(B)** Root-mean-square fluctuation (RMSF) by residue (calculated for the backbone atoms). Lilac-shaded bars indicate residues identified by TOPCONS Topology Analysis as being in the transmembrane α-helices. **(C)** STRIDE secondary structure analysis of the transmembrane domain simulated at 303 K. Alpha-helices, turns, 3-10 helices, and coils are indicated in pink, teal, blue, and white, respectively. Secondary structure is maintained throughout the 100 ns simulation (1001 frames).

**Supplementary Figure S3**

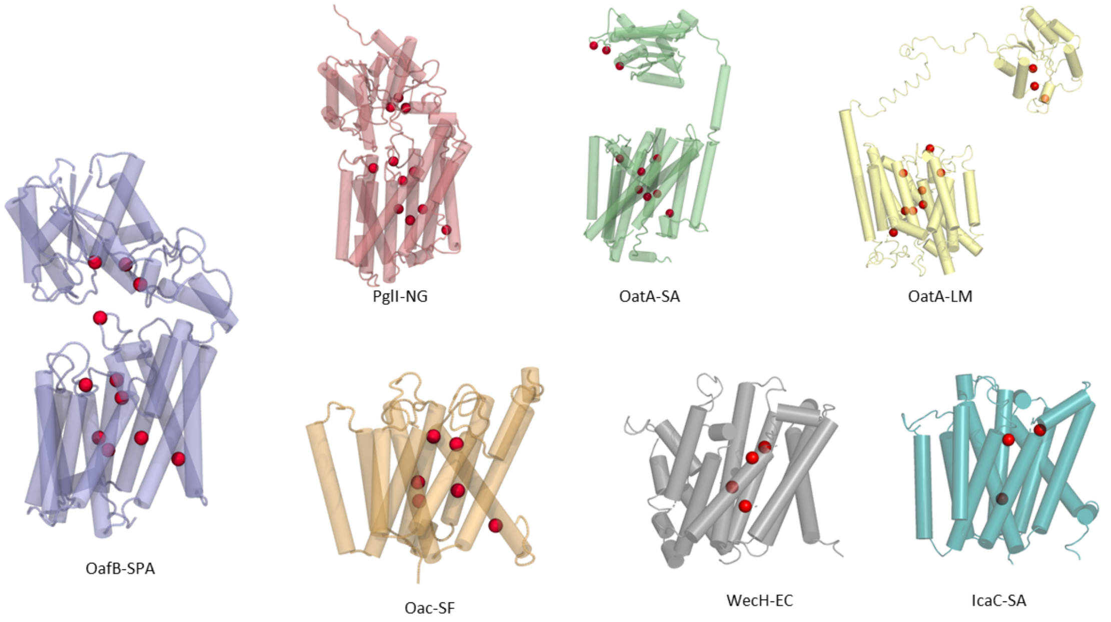


AT3 domain-containing proteins, with conserved essential residues (identified via sequence alignment) mapped onto their respective RaptorX structure predictions (red spheres). In all cases, these residues (excluding those in the catalytic triad of the SGNH domain) line a channel between helices within in the AT3 domain. This suggests that this family of proteins may have a common acyl donor and mechanism of action.

**Supplementary Figure S4**

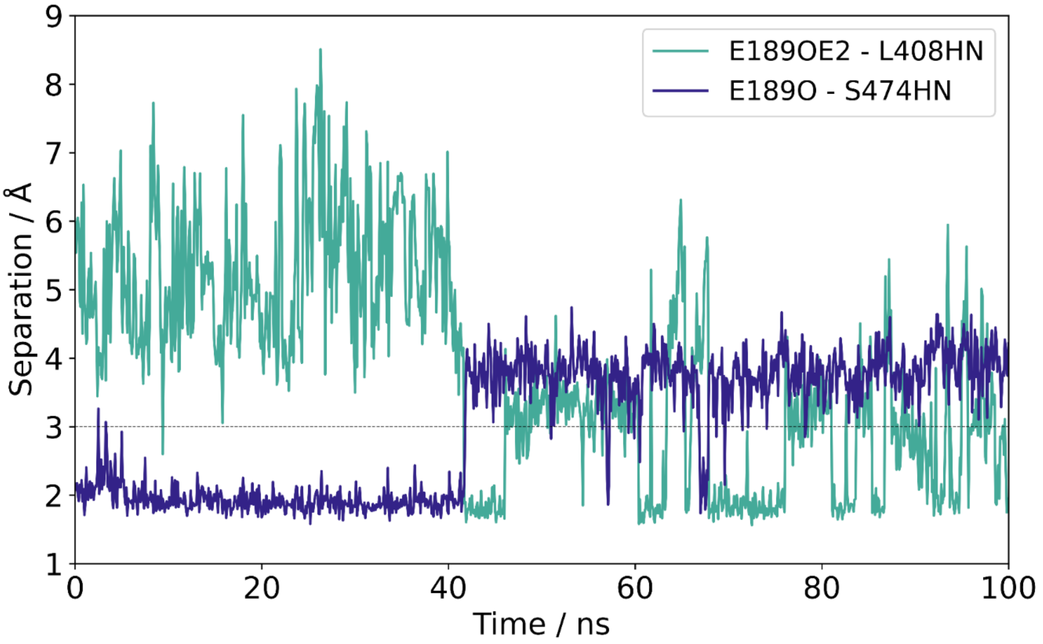


Separation between the carboxylate of E189 and backbone NH of L408 (teal), and backbone O of E189 and backbone NH of S474 (indigo). Cut-off distance (3 Angstroms) for hydrogen bonding shown as a dashed grey line. For the first 40 ns, the backbone carbonyl of E189 can form a hydrogen bond to the backbone amide NH of L474. After this point, the flexible loop in which E189 resides (between TMH5 and 6) shifts, breaking this interaction and allowing the formation of a hydrogen bond between the sidechain carboxylate of E189 and the backbone amide NH of L408 instead.

**Supplementary Figure S5**

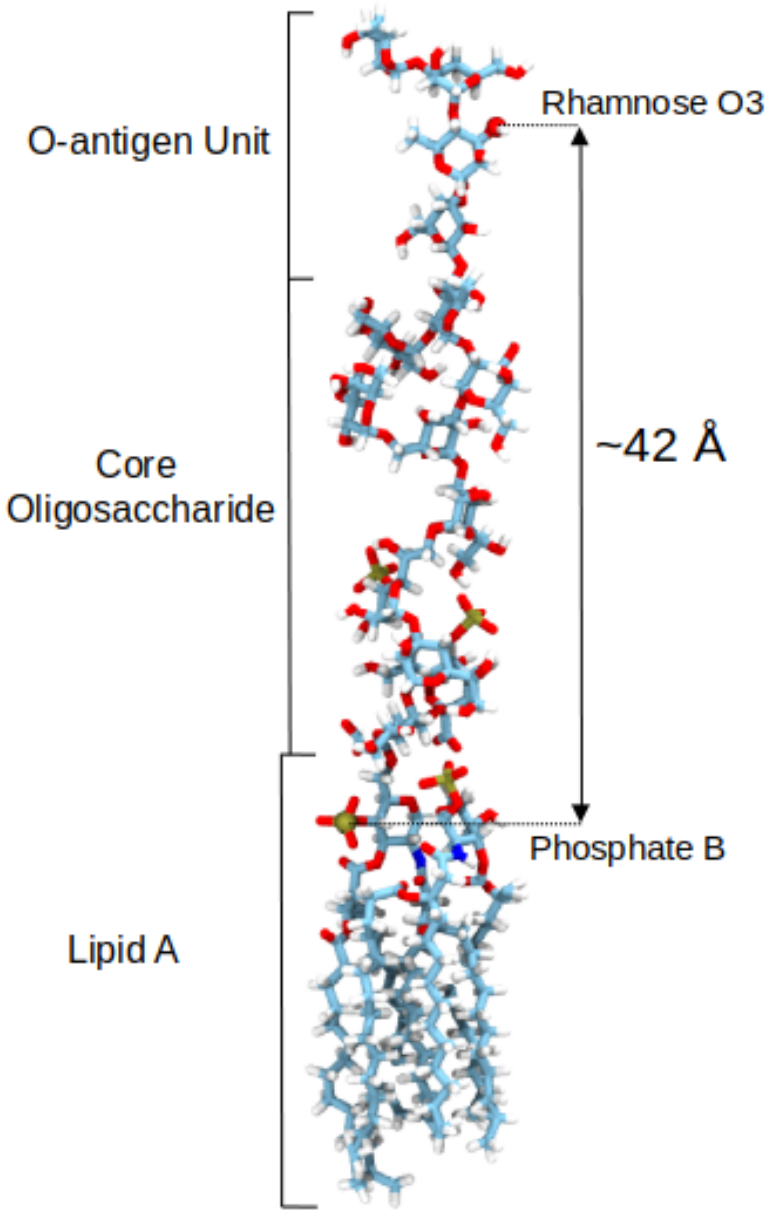


One molecule of LPS with a single O-antigen unit from *Salmonella enterica* subsp. *enterica* ser. Paratyphi A, shown in the Licorice representation. The acetylation site (O3 of rhamnose) and one phosphorus of the Lipid A are shown as spheres. This model was generated using the CHARMM-GUI LPS Modeller and shows the LPS unit before it has been embedded in the membrane and equilibrated. The separation between the Lipid A phosphate groups (which would be level with the phospholipid headgroups of the periplasmic leaflet of the inner membrane) are separated vertically from the acetylation site by a distance of 42 Å.
